# The selection arena in early human blastocysts resolves the pluripotent inner cell mass

**DOI:** 10.1101/318329

**Authors:** Manvendra Singh, Thomas J. Widmann, Vikas Bansal, Jose L. Cortes, Gerald G. Schumann, Stephanie Wunderlich, Ulrich Martin, Marta Garcia-Canadas, Jose L. Garcia-Perez, Laurence D. Hurst, Zsuzsanna Izsvák

## Abstract

Is the human early embryo unique in lacking an inner cell-mass (ICM) and having parallel development? We reanalyse single-cell transcriptomic data and stain human embryos *in situ* to reveal both classical step-wise development and a transcriptomically homologous ICM. This apparent classicism obscures phylogenetic singularity: unlike mice, human epiblast has self-renewal hallmarks and we have abundant blastocyst non-committed cells (NCCs), part of an apoptosis-mediated purging process. The transcriptomes of the pluripotent cells are fast evolving, in large part owing to endogenous retrovirus H (ERVH) activity, rendering all primate embryos unique. Each species is characterised by the ERVHs that are active and the neighbour genes whose expression are modulated. ERVH is associated with recent major gene expression gain and loss events of pluripotency­associated genes. Not least through lack of HERVH expression, the current portfolio of naïve cultures, putative *in vitro* mimics of pluripotent cells, are both developmentally and phylogenetically “confused”.

- Analysis of single cell transcriptomics and *in situ* stainings uncover, and enable characterization of, human inner cell mass (ICM)
- Cell purging via apoptosis defines a phylogenetically restricted class of blastocyst non-committed cells (NCCs), whereas HERVs in conjunction with host defence mark the committed cells of ICM
- Fast transcriptome evolution is particular to the pluripotent epiblast and is mostly due to the primate-specific transposable element, HERVH
- Current naive cultures don’t reflect human uniqueness being phylogenetically and developmentally “confused”.

## Introduction

While much that we know about early mammalian development is derived from knowledge in mice, in at least three aspects the human early embryo is thought to be different. First, while murine embryonic gene activation (EGA) occurs in the zygote (Flach et al., 1982), in primates it is later (at the 4/8 cells stage) (Braude et al., 1988). Second, in early human development an inner cell mass (ICM) is not evidenced and hence may not exist as a distinct lineage (Petropoulos et al., 2016; Sahakyan and Plath, 2016). While recent analyses (Boroviak et al., 2018; Stirparo et al., 2018) claim that ICM might yet exist, the cells identified express both ICM markers and apoptotic factors, suggesting a confusion of cell types. Third, the segregation of morula into trophectoderm (TE), epiblast (EPI) and primitive endoderm (PE) is thought to occur simultaneously rather than in the step-wise manner seen in mice and macaques (Chazaud and Yamanaka, 2016; Niakan et al., 2012; Sahakyan and Plath, 2016).

Despite the consensus that human pre-implantation embryogenesis is exceptional (Fuentes et al., 2018; Gao et al., 2018; Wu et al., 2018), even analysing single cell data (Blakeley et al., 2015; Petropoulos et al., 2016; Stirparo et al., 2018; Yan et al., 2013), the cells of the blastocyst still resist unambiguous identification. Consequently, while a definitive high-resolution reference transcriptome of all the cell types would be of great utility for the development of stem cell based regenerative therapies, such a description is lacking. Owing to the rapid progression of the developmental process, the most challenging task is to catch the distinct lineages during blastocyst formation. To address this, and to enable characterization of the stages of early human development, we use a plethora of approaches applied to single cell transcriptomic data both from humans and other primates. First, we employ a strategy of clustering both cell types and gene expression profiles using a dimension-reduction clustering methodology (Satija et al., 2015) that focuses on defining local resemblances. This strategy is well suited to single cell data, resolving transitory stages, capturing unresolvable clusters and, in turn, to identifying diagnostic markers, hence informing the biology of discrete cell types. Further, to add a time dimension and to provide lineage modelling, we additionally employ trajectory analysis, diffusion maps (DM) and diffusion pseudotime (DPT) to depict the transcriptional landscape of delineating cell types (Angerer et al., 2016; Haghverdi et al., 2016; Trapnell et al., 2014). For confirmation of key conclusions, we provide confocal immunofluorescent staining of early human embryos.

ICM in non-human species is a progenitor cell population, differentiating into EPI and PE developmental lineages in the blastocyst (Chazaud and Yamanaka, 2016; Nakamura et al., 2016). The expected hallmark of ICM cells is that such cells co-express multiple markers that are otherwise specific to EPI and PE (Chazaud et al., 2006). Our strategy identifies such cells in the expected location in the developmental trajectory. To confirm their classification, we compare the expression profile of these cells to that of known ICM cells of macaque. In addition, we provide stainings of human embryos to demonstrate the co-expression of multiple lineage defining genes in the single cell. We also find that the development process accords with the classical step-wise model.

These results allow us to characterize the transcriptome of progenitor ICM that gives rise to EPI and PE, suggesting that the human early embryo is not the outlier it was thought to be. In addition, we also resolve a previously unrecognised class of cells arising from morula that express no lineage markers but instead appear to be subjected to apoptosis. These we term non-committed cells (NCCs). Ontogenetic lineage analysis indicates that NCCs are excluded from the developmental process. Failure to recognize these has probably misled prior analyses (Stirparo et al., 2018), with, as a consequence, mischaracterization of putative progenitor ICM cells and the derivative EPI and PE lineages.

Given some level of fundamental homology between cell types across the mammals we can then ask whether the transcriptomes of the cell types evolve at the same rates and, coupled to this, whether any of our cell types appear to have phylogenetically unusual characteristics. The transcriptomes of the resolved cell types of the human early embryo are not equal in their rate of evolution, with the pluripotent ICM/EPI (mostly EPI) being highly divergent between primates while the transcriptomes of PE and TE remain relatively unchanged. Human EPI is unusual in having the transcriptomic hallmarks of self-renewal, not seen in mouse.

While we don’t approach the question of why the pluripotent cell transcriptomes might be uniquely fast evolving, to decipher the mechanism of accelerated evolution of EPI, we employ cultured pluripotent cells (PSCs) as a model for EPI. We follow the evolution of pluripotency by examining gene expression evolution between old-world (OWMs) and new world monkeys (NWMs) in primate PSCs, from human, bonobo, gorilla and marmoset. While, the OWMs share a high degree of similarity with humans in their genome sequence (92.5% to 97.5%) (Olson and Varki, 2003; Yan et al., 2013), the presence/absence differences are in no small part attributed to transposable element (TrE) insertions (Ramsay et al., 2017). Multiple waves of retroviral invasions into primate genomes, resulting in endogenous retroviruses (ERVs), are particularly important. After EGA, transcription of TrE families of different phylogenetic ages is evidenced and has a characteristic patterning (Friedli and Trono, 2015; Goke et al., 2015; Grow et al., 2015; Guo et al., 2014; Izsvak et al., 2016; Rowe and Trono, 2011; Smith et al., 2014). To decipher if (and how) TrEs contributed to transcriptome evolution during pre-implantation development within primates, in addition to considering classical genes as markers, we analyse transcripts associated with TrEs both in terms of changes to gene anatomy and in terms of expression dynamics. We highlight endogenous retrovirus H (ERVH) as a particularly key player in pluripotency transcriptome turnover and hence in defining human uniqueness.

The newly derived classification of early cell types and markers enables us to ask whether there is a pluripotent cell type in the human blastocyst that would be a good candidate for extraction and stable maintenance *in vitro* or to mimic. EPI having the hallmarks of self-renewal we highlight as the best candidate. Further, we can ask, given the unique nature of the pluripotent cells of the human embryo, whether any putative *in vitro* mimics, in the form of recently derived naïve cultures, accord both developmentally and phylogenetically in their transcriptomes with human *in vivo* pluripotent cells. The current portfolio of *in vitro* naïve cultures are heterogenous and even irreversibly modified (Pastor et al., 2016). We find them both developmentally and phylogenetically “confused”, and suggest experiments that might improve matters.

## Results

### Resolving the identity of cells expressing multiple and no lineage markers unmasks the human inner cell mass

What cell-types are present in the human pre-implantation embryo? In particular, is the inner cell mass really missing from the human blastocyst? To classify cell-types we used available single cell transcriptome data (Petropoulos et al., 2016; Yan et al., 2013). In contrast to previous analyses that identified differentially expressed genes (DEGs), we employ a strategy of clustering Most Variable Genes (MVGs) from whole cell transcriptomes, using a combination of clustering K-means and principal components (PCs) (Satija et al., 2015). We identified 1597 genes exhibiting high variability across single cells (on given thresholds) and thus potentially useful for defining cell types (Figures 1A and S1A). To reduce the dimensionality of the data we performed principle component analysis (PCA) and enlisted the significant PCs using a ‘jackstraw’ method (Satija et al., 2015). This identified nine significant principal components (p-value < 10^−10^). We used these PC loadings as inputs to t-distributed stochastic neighbour embedding (tSNE) for visualization. This approach allowed us to distinguish 10 clusters. These we classify on the basis of previously reported transcriptome markers (Petropoulos et al., 2016) (Figures 1A and S1A).

**Figure 1.**
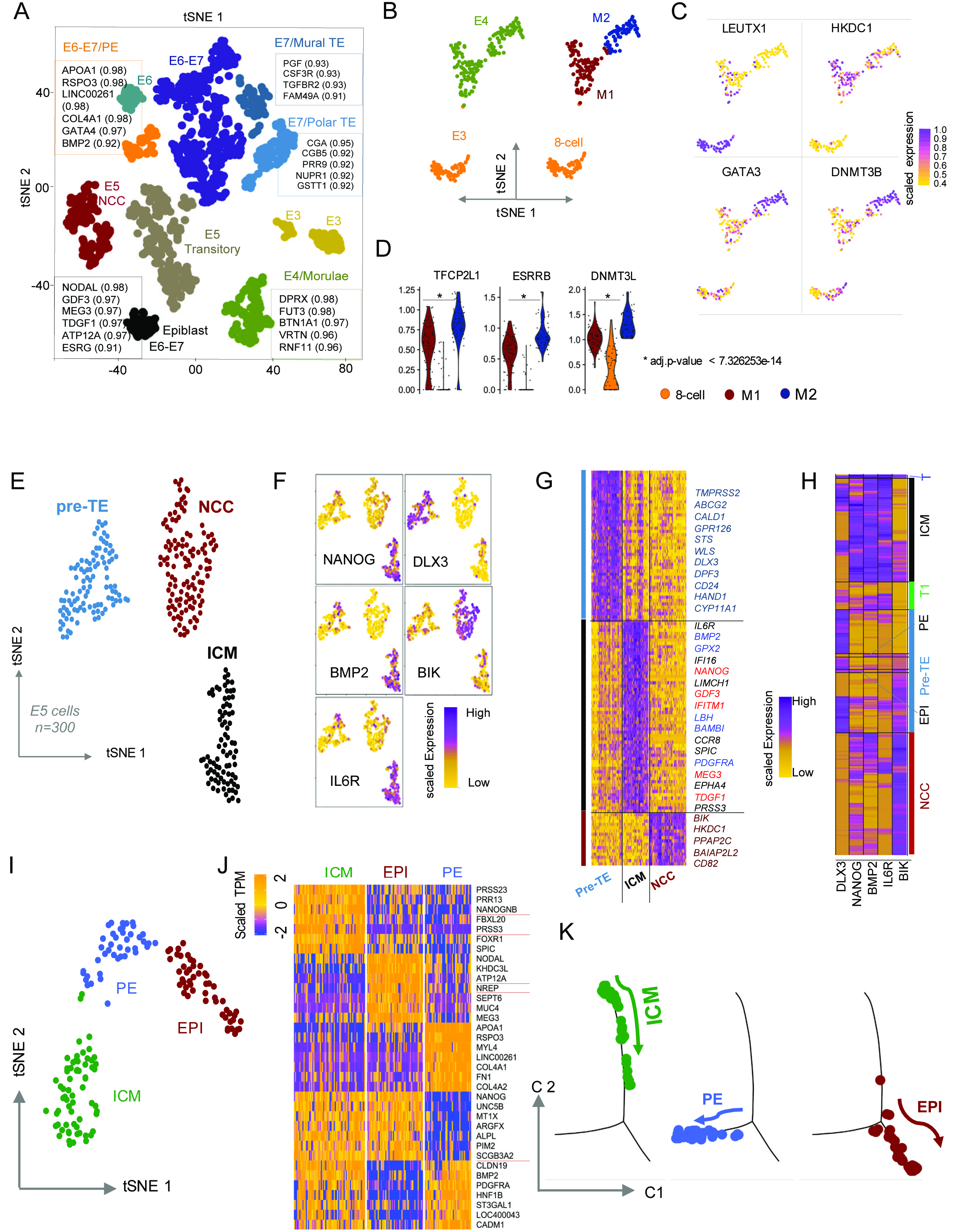
High-resolution dissection of human pre-implantation development. A. Two-dimensional t-SNE analysis of human single-cell pre-implantation transcriptomes using 1651 most variable genes (MVGs) resolves the following distinct cell populations: 8-cell at E3, morula at E4, pluripotent epiblast (EPI) at E6-E7, primitive endoderm (PE) at E6-E7, mural and polar trophoectoderm (TE) at E7. At E5, cells presenting none of the known lineage markers are referred to as non-committed cells (NCCs), whereas cells expressing multiple markers are annotated as transitory cells. The most discriminatory genes of each clusters are listed in boxes. Numbers in brackets refer to AUC values. Colours indicate unbiased classification via graph-based clustering, where each dot represents a single cell. B. tSNE unbiased clustering of E3-E4 cells using ~ 500 MVGs reveals three distinct cell populations (left panel). The three groups of cells are identified as 8 cells stage and two distinct populations of E4 blastomere (shown in dark red and midnight blue) named as Morula 1 (M1) and Morula 2 (M2) (right panel). Each dot represents a single cell. C. Feature plots based on tSNE plot from Figure 1B demonstrating lineage-specific expression of LEUTX (8-cell marker (Jouhilahti et al., 2016; Petropoulos et al., 2016; Yan et al., 2013), HKDC1 (M1 marker, this study), GATA3 (M2 marker, this study, but TE marker in (Blakeley et al., 2015; Petropoulos et al., 2016; Stirparo et al., 2018)} and DNMT3B (M2 marker, this study, but blastocyst marker in (Blakeley et al., 2015; Stirparo et al., 2018)). D. Violin plots showing the expression distribution of naive markers, suggested by human *in vitro* studies (Pastor et al., 2016; Takashima et al., 2014; Theunissen et al., 2016). TFCP2L1, ESRRB and DNMT3L in 8 cell and morula, M1 and M2 stages. Adjusted p-values are obtained by Benjamini-Hochberg (BH) correction. Each point represents a single cell. E. tSNE clustering of E5 cells using around 1000 MVGs reveals three distinct clusters of pre-TE, ICM, and a previously undocumented cluster of non-committed cells (NCCs). Each dot represents an individual cell. F. Feature plots based on t-SNE plot from Figure 1E visualising the expression of selected lineage-specific markers e.g. IL6R (ICM), NANOG (ICM/EPI), BMP2 (ICM/PE), DLX3 (pre-TE), BIK (NCC). Colour intensity gradient indicates the expression of the marker gene. Each dot represents an individual cell. G. Heatmap visualization of scaled expression [log TPM (transcripts per million)] values of distinctive set of 235 genes (AUC cut-off >0.90) for each cluster shown in Figure 1E (AUC cut-off >0.90). Colour scheme is based on z-score distribution from −2.5 (gold) to 2.5 (purple). Left margin colour bars highlight gene sets specific to the respective clusters. The ICM specific gene names in “red” or “blue” are progenitors and are also expressed at E6-E7 in EPI or PE cells, respectively. H. Heatmap of the row-wise scaled expression (log TPM) levels of selected marker genes for pre-TE (DLX3), pre-EPI (NANOG), pre-PE (BMP2), ICM (IL6R) and NCC (BIK). Colour bars next to heatmap were set manually showing the distinct group of cells expressing differential combination of markers. Note that transitory cells (T and T1) are co-expressing multiple lineage markers. I. tSNE biplot visualizes human ICM (*n*=71), PE (*n*=45) and EPI (*n*=52) clusters using the most variable genes (*n*=532) (See Table S3 for a full list of markers). J. Heatmap showing scaled expression (log TPM values) of distinctive marker gene sets defining EPI, PE and ICM. Genes specific to ICM include NANOGNB, PRSS3 and SPIC. Note the progenitor markers that are homogeneously expressed in ICM, but also in EPI (e.g. NANOG, MT1X) or PE (e.g. BMP2, PDGFRA). Note: Underlined genes are specific to human when compared with macaque. Colour scheme is based on z-score distribution, from –2.5 (gold) to 2.5 (purple). K. Monocle2 single-cell trajectory analysis and cell ordering along an artificial temporal continuum using the top 500 differentially expressed genes between ICM, EPI and PE populations. The transcriptome from each single cell represents a pseudotime point along an artificial time vector that denotes the progression of ICM to EPI and PE. Note that the artificial time point progression agrees with the biological time points (ICM is E5 which progresses to E6-E7 EPI and PE). For clarity, we show this trajectory in three facets.

In E1-E4, it is relatively straightforward to identify the known clusters corresponding to oocytes, zygote, 2, 4, 8 cells stage (E3) and even the more heterogeneous morula (E4) (Figure S1B). Morula, however, forms two subgroups that we term M1 (early) and M2 (late) (Figure 1B). While LEUTX1 flags the 8-cell stage, the two clusters of morula are marked by HKDC1 and GATA3, respectively (Figure 1C). Known blastocyst signature genes (e.g. TFCP2L1/LBP9, ESRRB, DNMT3L) are expressed in all morula cells, but more predominantly in M2 (Figure 1D).

Human blastocyst formation initiates at E5, progresses at E6 and stabilizes at E7 prior to implantation (Figure S1B). At E5, our tSNE plot visualizes expected and also previously unidentified clusters (Figure 1A). Our strategy identifies EPI and PE, as well as TE (polar, mural) clusters in E6 and E7 stages with their known markers (Petropoulos et al., 2016)[area under curve (AUC) ≥ 0.90] (Figures 1A, S1C and Table S1). A remaining cluster from E6 and one from E6-E7 are undefined since they express markers heterogeneously (Figures 1A and S1A).

Not all the clusters are as easy to classify as EPI, PE and the early cell types. Given heterogeneity in a cluster, might some cells also be transitory types? At E5, we observed two types of cell, either expressing multiple lineage markers or those that fail to express any markers of known blastocyst lineages (EPI-PE-TE: see Figure 1A). We find that the cells of the blastocyst that express multiple markers at E5 do not segregate on the tSNE plot, consistent with a reported transitory status (Petropoulos et al., 2016)(Figure 1A). By contrast, cells expressing none of the known markers form a distinct cell population. Given that these cells express no known lineage markers, we name them “non-committed cells” (NCCs) (Figures 1A and S1A).

If ICM exists, we would expect it to be resolved as a cell type segregating from morula. In order to further resolve the broad spectrum of transcriptomes of cells segregating from morula, we restricted analysis to E5 only, and removed cells with low quality transcriptomes (expressing (log2 Transcript Per Million (TPM < 1) ~ 5000 genes). This approach allows us to distinguish three clusters on tSNE (Figure 1E). Expression of DLX3, a known marker of a precursor population of TE defines the first cluster as pre-TE (Figures 1F-G). The second cluster, corresponding to the newly identified NCC, which, while expressing no lineage markers, homogeneously expresses apoptosis factor BIK (Figures 1F-G). Importantly, the third cluster while expressing specific markers (e.g. IL6R, SPIC), co-expresses known EPI (e.g. NANOG) and PE (e.g. BMP2) markers thus according with the definition of ICM (Figures 1F-G). Altogether, we identified 235 marker genes (AUC > 0.80) that are efficient molecular signatures of the individual clusters at E5 blastocyst (Figure 1G and Table S2).

### Human pre-implantation embryogenesis is a sequential process segregating from morula

Given identification of human ICM, we now ask whether the human progenitor ICM gives rise to EPI and PE, as this would challenge the recent ‘simultaneous model’ for human blastocyst formation. In a deviation from the sequential/step-wise lineage specification dynamics of mouse or macaque (Chazaud and Yamanaka, 2016; Nakamura et al., 2016), the new model suggests that the human morula segregates to EPI, PE and TE simultaneously around E5 (Petropoulos et al., 2016).

To evaluate the ‘simultaneous’ vs ‘sequential’ models, first we projected the top 100 most variable genes from the E5 blastocyst on diffusion maps. This approach resulted in a wishbone shaped pseudo-time trajectory (Figures S1D-E). We then mapped the scaled expression of our cluster-specific markers, DLX3 (pre-TE), BMP2 (PE), NANOG (EPI), IL6R (ICM) and BIK (NCC), on the diffusion components and determined co-expression patterns at single cell resolution. These strategies helped us to identify the transitory cells and decipher the developmental pathway following morula. Of 300 cells, we detected 3 expressing all four markers, indicating that these cells could be the precursors of early blastocysts. Another lineage of cells expressed the markers of the three layers of the blastocyst (n=25), featuring a transitory/precursor state (T1) prior the segregation to ICM (n=71) and pre-TE (n=86). ICM and either PE or EPI markers were enriched exclusively in the transitory cell population of T2 (n=8) and T3 (n=10), respectively (Figures 1H and S1D-E). We hypothesized that these few ICM-derived cells are the pre-EPI and pre-PE cells that would commit to PE and EPI at E6-7 stages. The identification of the transitory cells and ICM supports the model that human early embryogenesis accords with the classical step-wise process, as seen in mouse and macaque.

Additionally, we can also ask whether ICM, EPI and PE cells can be discriminated on dimension-reduced plots. This is valuable not least because the transcriptional flags to identify these populations in the midst of heterogeneous pool of cells would be an asset to stem cell biologists. We could detect the three distinct clusters of ICM, EPI and PE with marker genes (Figure 1I and Table S3). Consistent with its progenitor status, ICM homogenously co-expresses pre-EPI (e.g. NANOG, ARGFX, and SCGB3A2) and pre-PE (e.g. BMP2, PDGFRA and HNF1B) markers, whereas a set of genes specifically mark EPI (e.g. NODAL, MEG3, LEFTY2) or PE (e.g. APOA1, RSPO3, COL4A1) (Figures 1J and S1F, S2B). SPIC, NANOGNB and FOXR1 are among the genes specifically expressed in ICM (Figure 1J). Importantly, our pseudotime trajectory agrees with the biological timing and the bifurcation process of ICM giving rise to EPI and PE (Figure 1K). Thus, we can track the transcriptome changes on the branching points and define a set of genes that will be markers in the trajectories. Gradual alteration of gene expression over the pseudotime trajectories visualizes gene expression kinetics of ICM differentiating to PE and EPI (Figure S1F).

### *In situ* staining and comparative evidence confirm ICM identification

Above we have provided evidence both that the human ICM exists and that it differentiates in a sequential manner to EPI and PE. Here we provide two further evidences that we have correctly identified the missing human ICM. First, to confirm the identification of ICM we performed *in situ* stainings of human embryos. To visualize the ICM *in situ*, we performed immunostainings on E5 human embryos using antibodies against NANOG (characteristic of pre-EPI/EPI) and BMP2 (characteristic of pre-PE/PE). We expect that ICM cells in E5early would express both lineage markers, but that later (E5mid) cells would express one or the other as the ICM differentiates. As expected, in E5early, most cells co-express NANOG and BMP2 suggesting lineage divergence has yet to occur, consistent with the identity of the ICM as a progenitor population (Figures 2A and S2A-B). In E5mid, by contrast, we can identify cells with low to no expression of NANOG, and high expression of BMP2 (marking PE), as well as NANOG positive cells with no BMP2 staining (Figure 2A) indicative of EPI and PE fate specification post ICM. NCC cells express neither NANOG nor BMP2.

**Figure 2.**
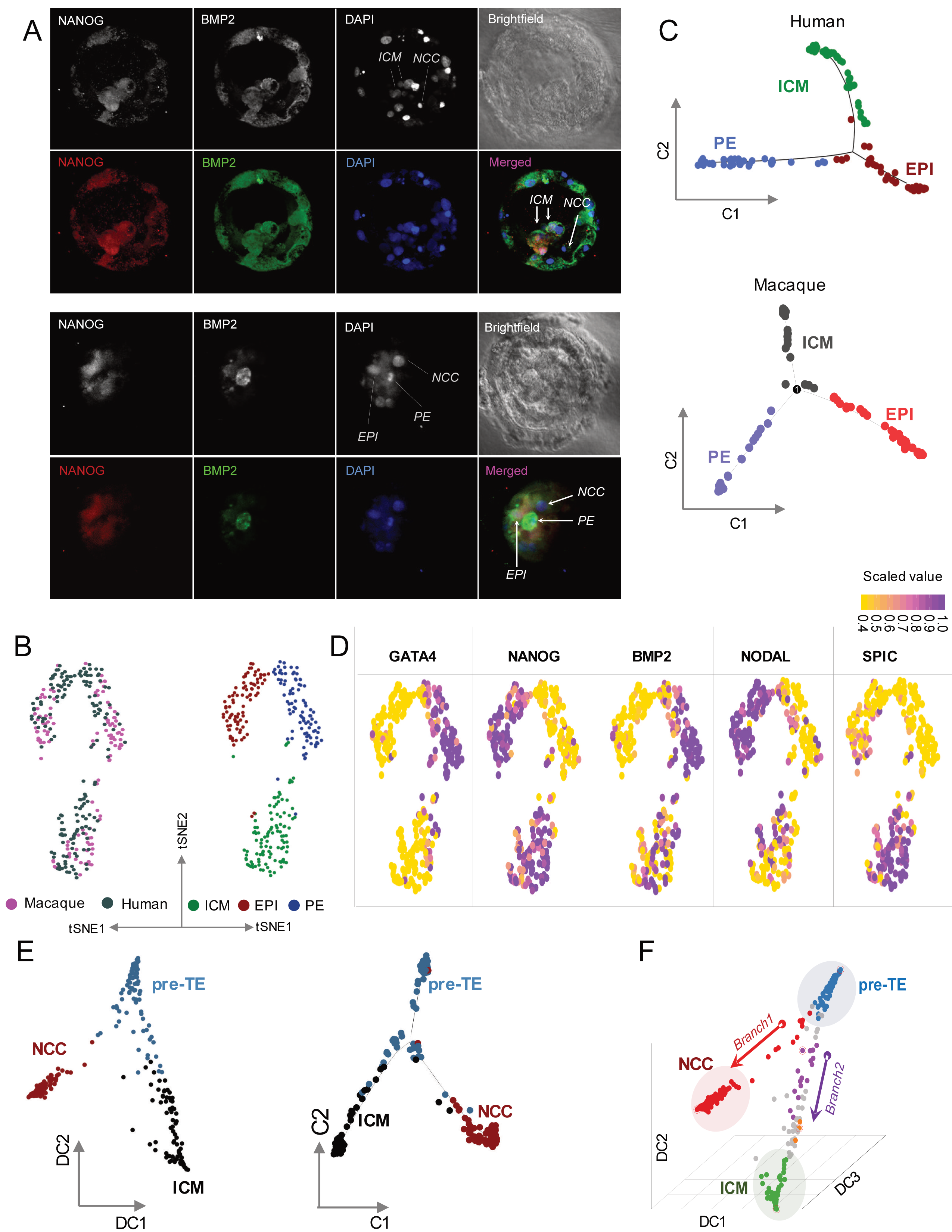
Immunostaining and comparative evidence confirm ICM identification. A. Representative confocal images (projections) of human E5early (upper two panels) and E5mid (lower two panels) blastocysts stained against NANOG (red), BMP2 (green) and DAPI, nuclear (blue) (bright field channel (black and white). Note that in E5arly embryos, NANOG and BMP2 co-stain cells of the ICM, whereas in E5mid embryos BMP2 and NANOG stained cells segregate, demonstrating the split of ICM into EPI(NANOG^+)^ and PE(BMP2^+)^ cells. Non-committed cells (NCCs) are expressing neither NANOG nor BMP2. B. tSNE plots of ICM, EPI and PE cells from *Homo* (169 cells, purple) and *Cynomolgus* (118 cells, grey) blastocysts after the normalization using “Seurat-alignment” (left panel). The joint clustering detects three distinct cross-species populations that can be identified as ICM, EPI and PE (right panel). C. Monocle2 pseudotime visualization of *Homo* and *Cynomolgus* ICM, EPI and PE trajectories using top ~500 differentially expressed genes. Note the similar dynamics of ICM differentiation to EPI and PE in the comparator species using orthologous genes. D. Unsupervised identification of shared lineage markers between *Homo* and *Cynomolgus*. Feature plots illustrate conserved gene expression in ICM, PE and EPI; SPIC (ICM), NANOG (ICM/EPI) and BMP2 (ICM/PE), NODAL (EPI) and GATA4 (PE) (the plots are in reference with Figure 2B). See Table S4 for full list of conserved markers. E. Diffusion map (DM) of E5 cells using the top 100 MVGs in 300 single cells plotted between Diffusion Components DC1 and DC2 (DC1 and DC2). DM identifies three separate branches of pre-TE, ICM and NCC (left panel). Pseudotime trajectory showing the ordering of E5 cells. Monocle2 visualization of pre-TE, ICM and NCC trajectories using DDRTree algorithms. The top 2000 differentially expressed genes are projected into a two-dimensional space (right panel). Colour codes as in (Figure 2B). F. Diffusion Pseudo Time (DPT) ordering of DM shown on Figure 2E. 3-D DM is plotted on Diffusion Components DC1, DC2, and DC3. DPT identifies the putative branching points of NCC and ICM (red and purple).

Second, we employ comparative transcriptomics to test for ontogenetic homology to known primate ICMs. We compared single cell transcriptomes of the blastocysts from human and a macaque (*Cynomolgus fascicularis*) (Nakamura et al., 2016; Petropoulos et al., 2016). Importantly, in contrast to the human study, the macaque blastocyst cells were pre-sorted by lineage markers (e.g. ICM, EPI, PE/hypoblast and TE) prior to sequencing. This had two consequences. First as NCCs weren’t known about they could not be extracted. Second, ICM cells were isolated on extraction and pre-defined as ICM. Are then the human ICM cells that we have identified homologous to these?

To compose comparable data for more detailed cross-species analysis, we use a recent method to integrate single cell transcriptomic data across different conditions (Butler et al., 2018). For comparison, we only use genes annotated in both species. The normalized cross-species single cell transcriptomic data allows us to perform well controlled comparative analyses. The reclassification of the merged transcriptomes on tSNE plots reveals a similar pattern of distinct cell types in both macaque and human (Figure 2B). Developmental lineage modelling positions the macaque lineages at an homologous ontogenetic position in the trajectory to our defined human ICM, PE and EPI, suggesting a phylogenetic conservation of the lineage specification in the blastocyst (Figure 2C). We find that ICM is unambiguously marked by the expression of SPIC in both species (Figure 2D and Table S4). As a progenitor of EPI and PE, ICM expresses both NANOG and BMP2 in both species (Figure 2D).

### Are non-committed cells parts of a quality control mechanism filtering out damaged cells?

What are the newly discovered non-committed cells (NCC) cells? Do they have developmental future and how do they relate to ICM? Employing diffusion methodologies at E5 projects three distinct cell populations (Figure 2E) on developmental trajectories and follows their hierarchical branching toward both ICM and NCC (Figures 2E and S1D-E). Expression of marker genes increases gradually as cells progress towards either ICM or NCC (Figure S1E). By ordering the cells according to their progression through ‘pseudotime’ supports a similar segregation dynamic (Figures 2E and S1D-E). Importantly, the observed high diffusion distance of NCCs suggests that these cells do not follow the developmental trajectory of either ICM or pre-TE, and might be excluded from the lineage specification process. This assumption is further supported by pseudo-temporal ordering of E5 cells that tracks NCCs as a deviating cell population from ICM and pre-TE trajectories with distinct branching points on the trajectory (Figures 2E-F).

Deriving from morula, we also expect ICM and pre-TE to share similar temporal coordinates at the end of E5. Indeed, ordering our previously defined cells on pseudo-temporal projections reveals that ICM and pre-TE share the same coordinates on the trajectory, while by contrast, NCC tracks as a deviating post-morula cell population (Figure S2C).

Previously we identified two subpopulations within morula (M1 and M2). Does this imply an unexpected very early specification of some cell types? How might these two types relate to NCC definition? Trajectory analysis including E3 to E5 stages reveals that morula-derived HKDC1 marked M1 cells eventually accumulate in NCCs, while M2 GATA3 positive cells are further trackable towards pre-TE, indicating that pre-TE is already programmed in the morula (Figures S2D-E). The cells of ICM, clearly separated from NCCs, are projected with either transitory cells or pre-TE (Figure S2C).

To discern more of the biology of NCCs, we determine the markers defining them. The top marker of NCCs, BIK (BCL2-Interacting Killer) (Figures 1E-H and S1E), is an apoptosis-inducing factor, suggesting again that this cell population has no developmental future. In addition to BIK, we observed the differential upregulation of numerous genes associated with programmed cell death (e.g. BAK1, various caspases or MAPK3, etc.) (Figure S2F).

Why might the NCCs be expressing apoptotic factors? This may be a mechanism that serves as a quality control measure to eradicate damaged cells or that simply removes surplus cells during the developmental process. In the former context, one strong possibility is that they might be damaged by the activity of mutagenic transposable elements (TrEs). In humans, the phylogenetically young elements include certain transpositionally active, mutagenic TrEs, such as Long Interspersed Element class 1 (LINE-1 or L1), SVA and Alu (Hancks and Kazazian, 2012; Mills et al., 2007). Importantly, the majority of the mutagenic elements in the human genome are activated following EGA with their expression peaking in morula (Goke et al., 2015; Romer et al., 2017). Thus, we would expect that activation of mutagenic TrEs could adversely affect some cells in the embryo. The quality control hypothesis thus predicts that NCCs should express higher level of young TrEs with transposing potential, while committed cells would repress expression of such TrEs.

To monitor TrE expression dynamics during pre-implantation embryogenesis, we first analyse bulk RNAseq datasets (Wu et al., 2018). This approach reveals a sharply distinct expression pattern of TrEs, featured by top signals for the SVA_D/E and HERVH at the 8-cell stage and bulk ICM, respectively (Fig. S3A). However, bulk RNAseq appears not to be suitable to distinguish committed from non-committed cell populations, as we are readily detecting NCC markers (e.g. BIK) contaminating the ICM sample (Figure S3A). Thus, to compare TrE expression in NCCs against ICM, we analyse single cell RNAseq data. Using averaged expression (Log2 CPM+1) of each particular TrE family across the cell populations clustered as ICM and NCC at the diffusion map (see Figure 2F), we observe a transcriptional pattern of TrEs that distinguishes ICM from NCCs (Figure 3A). Curiously, while the activated families in the ICM include HERVH, we find SVA_D/E upregulation in NCCs. This analysis reveals a more general pattern, namely that transcriptionally activated TrEs in ICM are relatively phylogenetically old (> 7 MY) and, crucially, transpositionally inactive, whereas the upregulated TrEs in NCCs are young (< 7 MY) and include mutagenic elements (Figure 3A). In ICM, the abundant transcripts of various older, endogenous retroviruses (ERVs) are dominantly represented by their full-length versions: LTR2B/ERVL18, LTR41B/ERVE_a, LTR17/ERV17, LTR10/ERVI, MER48/ERVH48 and LTR7/(H)ERVH in ascending order of average expression. The list of activated young TrEs in NCCs includes both LTR-containing TrE such as LTR5_Hs and recently mutagenic non-LTR retrotransposons, such as AluY (Ya5), SVA_D/E and L1_Hs elements (Figure 3A).

**Figure 3.**
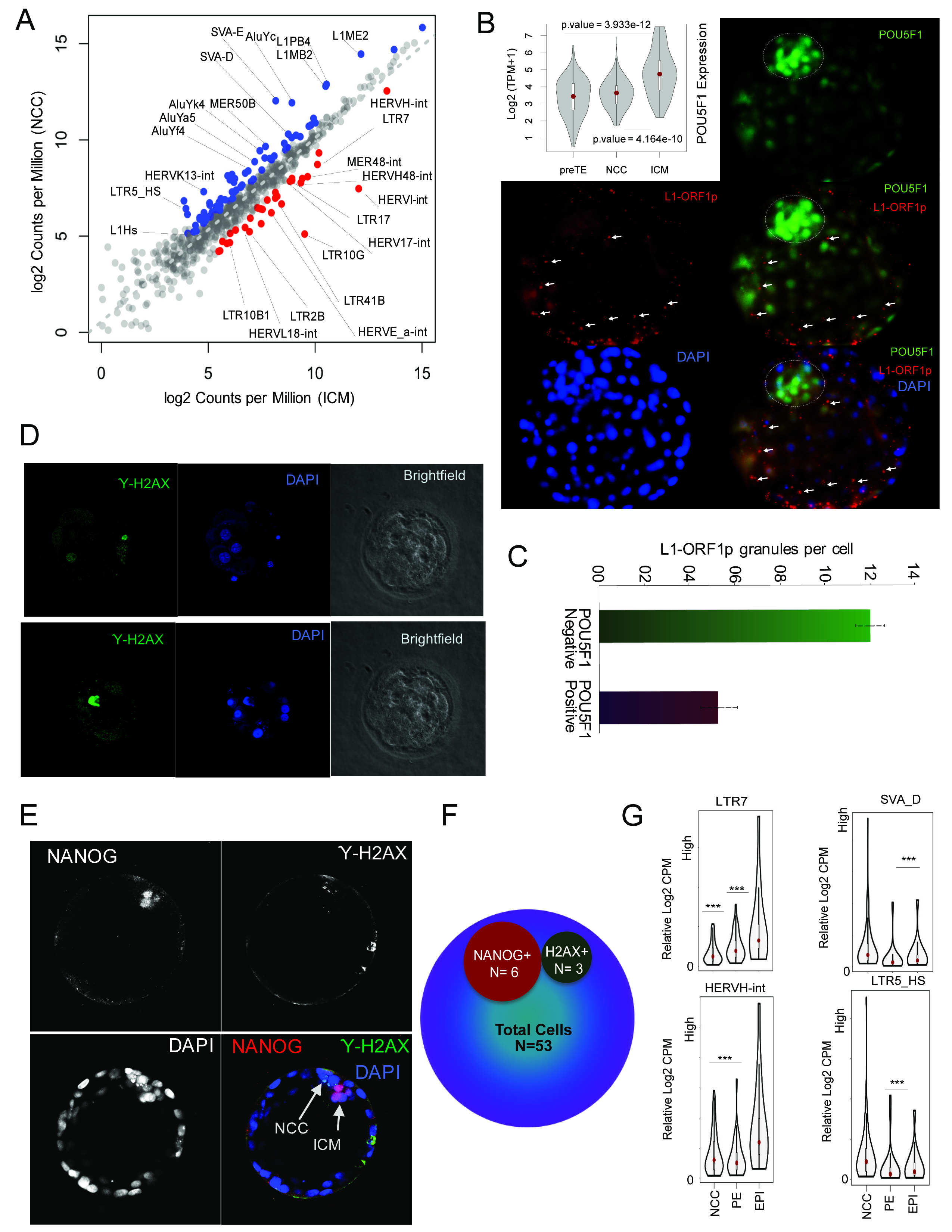
Characterization of the non-committed cell population in E5. A. Phylogenetically young (< 7MY) and old (> 7MY) transposable elements (TrEs) are antagonistically expressed in non-committed cells (NCCs) and the inner cell mass (ICM). The scatterplot shows the comparison of normalized mean expression in CPM (Counts Per Million) of various TrE families between the averaged pool of ICM (x-axis) and NCC (y-axis) cells. Read counts per TrE family are normalized to total mappable reads per million. Note: Uniquely mapped reads were considered as one alignment per read. Multimapping reads were considered as one alignment only if they were mapped to multiple loci, but exclusively within a TrE family. Every dot corresponds to a TrE family. TrE families enriched in ICM (red) vs NCC (blue). B. Representative confocal images show immunofluorescence staining in human early (E5) blastocysts with anti POU5F1/OCT4 (nuclear, green), L1-ORF1p (cytoplasmic granular, red), DAPI (nuclear, blue). Note: POU5F1 ^+^ cells are significantly enriched in the ICM (circled) and compacting near polar TE. A violin plot (upper left panel) visualizes the density and expressional dynamics of the POU5F1 in pre-TE, NCC and ICM at E5. Co-staining demonstrates the exclusive expression of POU5F1 and L1_ORF1p during the formation of blastocyst. The cells expressing higher POU5F1 compacting to form the ICM at the polar region of the blastocyst are less well stained for L1-ORF1p. L1-ORF1 stains scattered cells and pre-TE, not included in the compacted population of cells (arrows). L1 (LINE-1_Hs) belongs to a group of mutagenic, Young TrEs and supports transposition of both LINE-1 and the non-autonomous Alu and SVA elements. See also Movie 1. C. Numerical analyses of L1-ORF1p cytoplasmic foci of expression in cells that are POU5F1^−^ vs POU5F1 ^+^ at E5 blastocoel. The graph shows the average number of L1-ORF1p cytoplasmic foci in POU5F1^−^ and POU5F1 ^+^ cells, in the ICM and the blastocoel (pre-TE cells were not considered for this analysis), with standard deviation. D. Double stranded DNA damage visualized by H2AXγ immuno-staining is readily detected in a fraction of cells during blastocyst formation. Representative immunofluorescent H2AXγ staining (green), DAPI (blue) in two (upper and lower panels) human E5 blastocysts (brightfield, left panels). Damaged/dying cells accumulate H2AXγ signals. H2AXγ stained cells might still have integrity and exhibit normal oval shape (upper panel) or loose integrity (lower panel). Note: As cells might die upon thawing, we only analysed blastocysts that fully developed *in vitro* from E2 embryos. E. Representative confocal immunofluorescent images of human E5 blastocysts co-stained against NANOG (red), H2AXγ (green) and DAPI (blue); (brightfield, left three panels). Cells in the blastocoel are stained either and exclusively with NANOG, representing the compacting cells of the ICM, or with H2AXγ, representing damaged/dying cells. Note the disintegrating nucleus in H2AXγ^+^ cell (NCC). F. Venn diagram shows the schematic numerical analysis of immunofluorescent co-staining shown on Figure 3E. G. Violin plots visualize the density and antagonistic expressional dynamics of transposable elements, SVA_D, LTR5_HS (Young) vs LTR7/HERVH (Old) in non-committed (NCC) vs committed cell populations (PE and EPI) of the human blastocyst. Note that Young elements are enriched in NCCs vs committed cells (adjusted-p-value < 0.000072), whereas LTR7/HERVH-int are enriched in EPI vs PE and NCCs (adjusted-p-value < 0.00037).

To corroborate these findings, we analysed human pre-implantation embryos using confocal microscopy and an antibody raised against the L1_Hs-encoded L1_Hs-ORF1p protein (Macia et al., 2017). Our immunostaining detects robust expression of the L1_Hs-ORF1p during blastocyst formation, *in situ* (Figures 3B and S3B and Movie 1) with an inverse correlation with POUF5F1 (OCT4) levels (Figure 3C). The cells that show high intensity of POU5F1/OCT4 staining are compacting near polar TE to form ICM whereas, L1_HS-ORF1p stains scattered cells. Some of the excluded cells from the developing ICM may be progenitors of trophectoderm (Figures 3B and S3B) (see also Muñoz-Lopez. et al., 2018). However, we also detect cells with signs of genomic DNA damage, visualised by H2AXγ staining (Figure 3D). OCT4^low^ L1_HS H2AXγ^+^ cells, we presume, fail to express a commitment marker, consistent with NCCs having no future and being filtered out from the developmental process due to DNA damage (Gasior et al., 2006). Although, POU5F1/OCT4 is highly expressed in ICM, it is also detected in all E5 cells, thus we also use additional staining with NANOG that marks ICM/EPI more precisely (Figure 3E). Numerical analyses support that OCT4 /NANOG cells don’t have the DNA damage marked by H2AXγ (Figure 3F). While, large-scale transcriptional upregulation of TrEs might itself trigger apoptosis (Bouttier et al., 2016), we speculate that the DNA damage, generated by transposition (e.g. L1_Hs and SVA/Alu - L1-dependent non-autonomous elements) contributes to the apoptotic process in NCCs. The quality control hypothesis suggests that cells are selectively dying, and further predicts that NCCs are a developmental dead end. Consistent with the model, NCCs diffuse out from the developmental trajectory, are not detectable after E5, consistent with exclusion from the developmental process (Figures 1E-F, 2E-F and S1D-E, S2C-E).

### Host defence, in part owing to relatively old transposable elements, mark committed cells of human ICM

Given the potential mutagenicity of certain young TrEs, it is expected that young TrEs are efficiently silenced in the ICM (Figure 3A), but perhaps not entirely in the pre-TE (Figure 3B) (see also Muñoz-Lopez. et al., 2018). This accords with the expression of host defence genes such as IFITM1 and APOBEC3C/D/G (Figure S3C) suppressing viral/TrE activities (Grow et al., 2015; Knisbacher et al., 2016). The suppression of the younger TrEs that are active in NCC (i.e. SVA and LTR5_HS), is not peculiar to ICM but is also seen in committed cells derived from ICM, namely EPI and PE (Figure 3G). EPI is also enriched with LTR7/HERVH transcripts, these being at low levels in NCCs.

This pattern of mutual exclusion of Young and Old TrEs, the former being seen in NCCs, the latter in ICM, could be owing to some third-party switches. Alternatively, activity of one might suppress the activity of the other. As HERVH expression is expected to decline following implantation, knocking down (KD) HERVH in hPSCs can model certain aspects of this developmental stage, when cells discontinue to self-renew and commit to differentiate (Lu et al., 2014; Wang et al., 2014). The transcriptome of KD-HERVH_H1 cells (Lu et al., 2014) reveals upregulation of Young TrEs (Figure S3D), suggesting that the activation of ancient TrEs (e.g. HERVH) is associated with the maintenance of genome stability. Among the downregulated genes in KD-HERVH_H1, we identify APOBEC3G (Figure S3E), located downstream of a full-length HERVH. As HERVH can regulate its own transcription *in trans* (Lu et al., 2014; Wang et al., 2014), it can affect the expression of several HERVH loci, including its potential target genes *in cis* (e.g. APOBEC3C/D/G) (Figures S3F). In addition, we observe a relative upregulation of genes in the neighbourhood of active HERVHs (10 KB window) (Figure S3G), suggesting a potential *cis*-regulatory effect of HERVH on its neighbours. APOBEC3G, a member of the cytidine deaminase gene family, is implicated as a suppressor of young TrEs (including LINE-1) (Hulme et al., 2007; Khatua et al., 2010), indicating one mechanism by which HERVH could exercise control over genome stability.

### The transcriptome of EPI is fast evolving

As HERVH has been recently implicated in contributing to human pluripotency regulation (Lu et al., 2014; Wang et al., 2014), and the macaque genome carries relatively few copies of these full-length elements (~ 50), we asked if besides the obvious phylogenetic conservation, it was also possible to track the cross-species differences in pluripotent stem cells of the blastocyst. To determine differences between *Homo* and *Cynomolgus*, we analysed orthologous genes and identified the genes that are expressed in both of the species or that are species-specific in their expression. In human, we observe a significant expression (Log2 TPM > 2) of 30 genes in ICM and 20 in PE that are not expressed in *Cynomolgus* (Log2 TPM < 1 in < 5 % cells). By contrast, we see many more (48) EPI specific genes (expressed in > 95% cells) (Figures 4A-B and Table S5).

What underpins this greater divergence of EPI between macaque and human? To approach the problem of uniqueness more generally, we first ask whether the transcriptomes of the different cell types are evolving at different rates. A three-dimensional tSNE analysis of the merged transcriptome reveals that while the macaque/human ICM and PE transcriptomes are clustering together, the EPIs form distant sub-clusters (Figure S4A). The observed divergence is supported by trajectory building using defined criterion, as the PE and ICM transcriptomes follow a similar trajectory, whereas the macaque and human EPIs take a different path (Figure 4C), indicative of the transcriptome of the EPI being faster evolving than the other cell types in the blastocyst.

**Figure 4.**
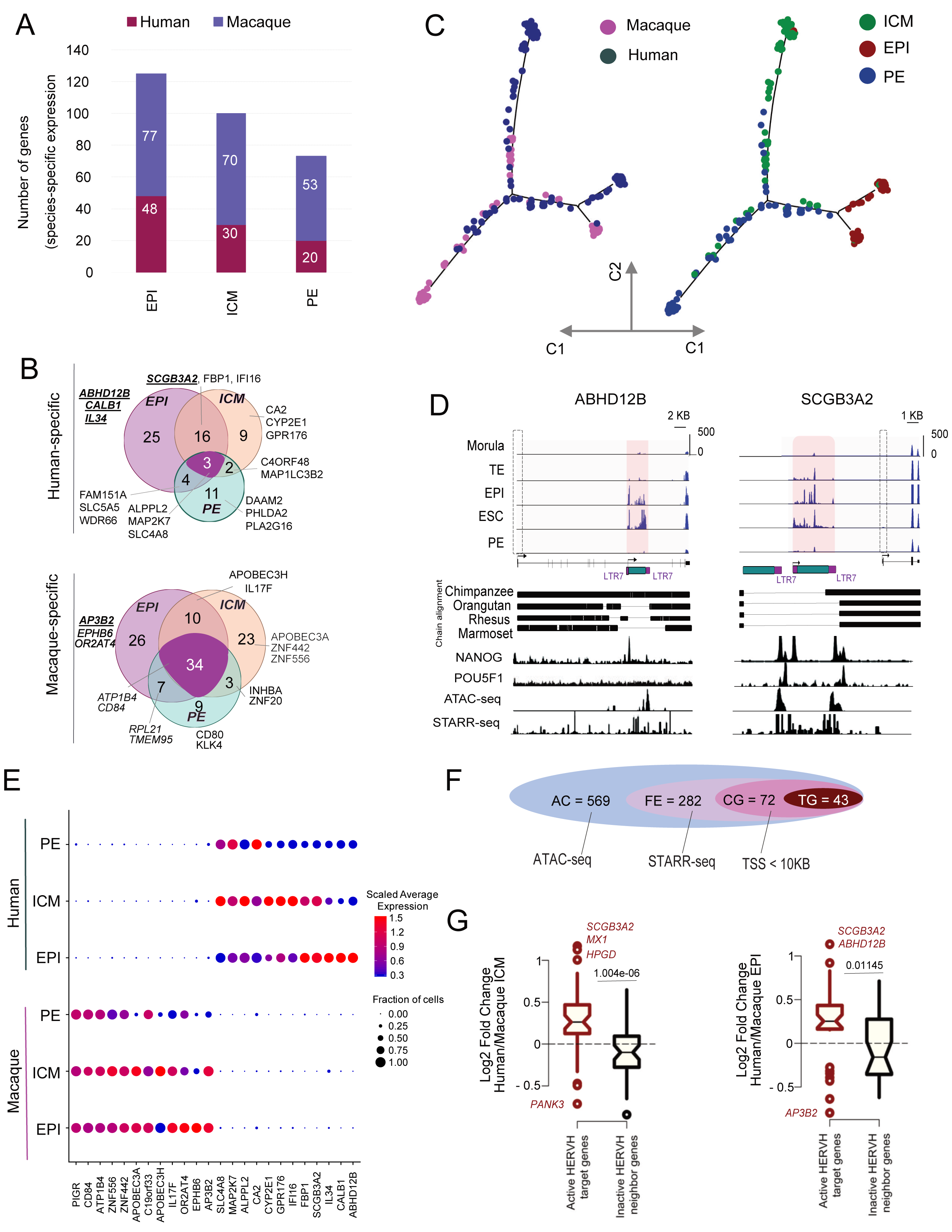
Accelerated divergence of the pluripotent epiblast during primate evolution. A. Stacked barplot shows the species-specific expression (log2 TPM > 1) of orthologous genes in EPI, ICM and PE lineages. Species-specific gene expression is considered to occur when the gene is expressed in at least 95% of the cells of a given lineage, but not in the same lineage (< 5% of the cells) of the other species. Note that species-specific gene expression is enriched in EPI vs ICM or PE. B. Venn diagrams visualizing species-specific gene expression (shown as numbers in previous figure) in the ICM, EPI, PE lineages of the blastocyst. Human (upper panel) vs macaque (lower panel) show depicted species-specific genes, which are significantly upregulated in a particular lineage (FDR < 0.05) or shared between lineages (e.g. have insignificant differential expression between the lineages). Genes shown in “italic” are located next to a HERVH in the human genome (10KB window). Genes shown in “bold” and “underlined” are HERVH target genes identified by the combined ATAC-seq and STARR-seq analyses in pluripotent stem cells (see also Figure 4D). Note the higher divergence of human blastocyst lineages. For full list of genes with species-specific expression see also Table S4. C. Pseudotime trajectory analysis of cross-species ICM, EPI and PE cell populations from human (N=169 cells) and macaque (N=118 cells) blastocysts merged in a single data frame (shown on Figure 2B). This reveals four branches, based on the 250 top DEGs (left panel). Annotation of the distinct lineages over the trajectory (right panel). Note that ICM is branching into a conserved human/macaque PE, while human and macaque EPIs are diverged (plotted against the first two components). D. HERVH-remodelled genes are incorporated into the regulation of human-specific pluripotency. HERVH-mediated remodelling of ABHD12B and SCGB3A2 genes. Integrative Genome Visualization (IGV) of uniquely mapped reads of single cell RNAseq (upper panel), ATAC-seq, STARRseq, NANOG (ChiPseq) (bottom) over ABHD12B and SCGB3A2 and their proximal full-length HERVH loci (LTR7-HERVH-int-LTR7). Arrows show the annotated and LTR7/HERVH-enforced transcriptional start sites (TSSs). Transcription skipped at annotated (empty dashed box) and HERVH-enforced TSSs (shaded box) are shown. Both genes lose their annotated TSS and proximal exons to form HERVH chimera. The chimeric transcripts exhibit a specific developmental pattern. The lowest panels show phylogenetic conservation status, the presence (thick line) and the absence (narrow line) of the human sequence compared to the *Chimpanzee*, *Orangutan*, *Rhesus* and *Marmoset* assemblies. While, HERVH at the ABHD12B locus appears to be intact in Chimpanzee, it has several deletions compared to the human version (white columns in the thick line). E. Dot plot showing the scaled expression dynamics of selected genes in species-specific pattern. The size of the dots is proportional to the fraction of cells expressing the given gene. Colours represent an average Log2 expression level scaled to number unique molecular identification (UMI) values in single cells. F. Schematic representation of the step-wise strategy used to identify HERVH target genes in human pluripotent stem cells (PSCs) (bulk ICM and *in vitro* hPSCs). AC: Accessible Chromatin at HERVH loci in bulk ICM (ATAC-seq, RPKM > 2); FE: Functional Enhancer HERVH (AC + STARR-seq, RPPM > 144); CG: Cis Genes (distance of Gene code TSS from FE, |10KB|); TG: Target Genes (No ATAC-seq peaks (FPKM > 2 per 100 bp) between FE and CG). G. Notched boxplot represents the distribution of average difference (at Log2 scale) of HERVH neighbour target gene expression is significantly enriched in human versus macaque ICM (left) and EPI (right) lineages. Cell populations are compared independently and in a pairwise manner. Only orthologous genes, annotated in both human and macaque Refseq gtf were analysed. Note: HERVH target genes can be upregulated (e.g. SCGB3A2) or tuned down (e.g. AP3B2) in the human vs macaque comparison. Certain HERVH-affected target gene expression is shared between ICM (left panel) and EPI (right panel), but could be also different.

To understand what might underlie this, we sought to classify the gene regulatory networks in the distinct blastocyst lineages. To investigate the gene regulatory networks (regulons), we applied the Single-Cell Regulatory Network Inference and Clustering (SCENIC) method (Aibar et al., 2017). This examines DEGs for master regulators and their putative target genes as regulons. SCENIC identified NODAL, GDF3, TDGF1/CRIPTO as the specific master regulators of EPI (Figure S4B-C and Table S7). Among the target genes of the EPI-specific master regulators we find several HERVH-driven/remodelled genes (Figure S4C). Importantly, these HERVH-affected genes (e.g. SCGB3A2, ABDH12B, IL34) exhibit human-specific expression (Figures 4D-E and S4E), pointing to their potential contribution to the evolution of EPI function. In order to understand the progression of evolutionary changes in regulating pluripotency, we employ pairwise weight correlation network analysis (WGCNA) on human-macaque orthologous gene expression (observed similar modules as in SCENIC analysis), and extract significant pairwise ranked correlations (>80%) on scaled data. Besides conserved gene expression (e.g. NODAL, GDF3, TDGF1 and PRDM14), this strategy identifies genes whose expression has been shifted between cell types in the two species (e.g. MT1G, MT1X from ICM in *Cynomolgus* to EPI in *Homo*; ATP12A, from TE in *Cynomolgus* to EPI in *Homo*), or are unique to Homo EPI (e.g. SCGB3A2, ABHD12B) (Figures 4D-E and S4A,D).

### Human-specific HERVH-remodelled genes modulate pluripotency

Among the human-specific transcripts identified above, we find several HERVH-remodelled genes, suggesting that expressed HERVHs are capable of enforcing chimeric transcription (Wang et al., 2014). For example, while ABHD12B and SCGB3A2 are both conserved in sequence in *Cynomolgus* and *Homo*, the gene body of the human version of each has been remodelled by HERVH and expressed exclusively in human pluripotent cells (Figure 4D). SCGB3A2, driven by both NANOG/POU5F1 shows expression in both ICM and EPI, whereas ABHD12B and other HERVH-derived transcripts (e.g. ESRG, HHLA1) are expressed more abundantly in EPI (Figures S4A and S4E-F). Conversely, while the non HERVH associated TFPI2 is exclusively expressed in EPI, the expression of its HERVH-remodelled paralogue, TFPI, occurs in both human ICM and EPI (Figures S4E-F). These examples are consistent with a functional diversification of HERVH-enforced gene regulation (or HERVH-associated transcripts) in modulating pluripotency. Supporting this assumption, a number of HERVH-derived transcripts and remodelled genes have been confirmed to be functional and to affect pluripotency (Durruthy-Durruthy et al., 2016; Loewer et al., 2010; Wang et al., 2014; Zhao et al., 2007).

### Transcriptional rewiring is enabled by LTR7/HERVH functioning as an enhancer

How might LTR7/HERVHs be acting to rewire transcriptional networks? One possibility for network structuring, supported by prior in-depth analyses (Lu et al., 2014; Wang et al., 2014), suggests that HERVH comes preloaded with transcription factor (TF) binding sites enabling activity as a potential promoter/enhancer. To identify functional enhancers regulating human pluripotency, we analysed genome-wide STARR-seq datasets from cultured PSCs (GSE99631) (Barakat et al., 2018) and ordered TrE-derived sequences according to their potential strength (Figure S4G). This strategy puts LTR7, LTR5_Hs and HSAT on the top of the list. Significantly, besides its LTR (LTR7), the internal HERVH sequence (HERVH-int) is also predicted to have a strong enhancer function. Considering their relatively large number, LTR7/HERVH-int are expected to have the most significant impact on human pluripotency (Figure S4G).

Can we more fully implicate HERVH as causative of the neighbour gene effects? To answer this, we add an additional layer by analysing expressed LTR7/HERVH loci in the dataset generated by ATAC-seq (GSE101571) (Barakat et al., 2018; Wu et al., 2018). This strategy identifies 569 HERVH loci at accessible chromatin regions in bulk ICM (ATAC-seq RPKM > 2), around half of which (N=282) are also functional enhancers in various pluripotent stem cells (PSCs). When an identified HERVH enhancer is the closest in cis to an actively expressed gene, we consider that gene to be a target (N=43) of a functional HERVH enhancer (Figure 4F).

We also asked whether human and macaque orthologous genes expressed in ICM or EPI are potentially HERVH-driven. As a control, we look at genes in the proximity of inactive HERVH (STARR-seq RPPM < 144). The genes in proximity to transcriptionally active HERVH are relatively active in human PSCs compared with macaque counterparts (where most of the ERVH loci are absent), consistent with HERVH promoting gene expression in the neighbourhood. This cross-species analysis identifies HERVH target genes whose expression is upregulated or tuned down in either ICM or EPI in human relative to macaque (Figure 4G). In addition, there are genes for which no macaque ortholog is found that have been remodelled/created by HERVH and expressed exclusively in human ICM and EPI (notably ESRG and LincROR) (Table S1).

### The divergence of the pluripotency transcriptome following the split of old and new world monkeys is owing to HERVH enforced expression

HERVH was introduced into the primate genome after the old world/new world monkey split. As HERVH-enforced gene regulation/remodelling appears to be involved in the evolution of EPI, we thus further dissected the emergence of the HERVH-driven regulatory network of pluripotency in primates. Our expectation is that much of the divergence in regulation will be owing to HERVH.

As models of the pluripotent EPI, we use comparable pluripotent stem cells (PSCs), established from human and various Non-Human Primates (NHPs). To determine differentially expressed genomic loci between human and NHPs transcriptomes, we include male PSCs from human, chimp, bonobo (Marchetto et al., 2013) and our own *Gorilla* data (Wunderlich et al., 2014). We additionally generate RNASeq data from *Callithrix* (Muller et al., 2009), where HERVH is not present (Izsvak et al., 2016) so as to better estimate the pre-ERVH ancestral state. We also extract HERVH-governed genes defined as those differentially regulated in the knockdown cells (HERVH-KD) in the human embryonic stem cell line ESC_H1 compared to a control (GFPcontrol-KD) (Lu et al., 2014). We processed and normalized the datasets to perform various comparative analyses (Figures S5A-D).

Our cross-species mapping demonstrates how dramatically the expression of human EPI markers (e.g. including LEFTY1/2, NODAL) changes between human and *Callithrix* PSCs (Figure 5A), supporting the large-scale restructuring of the pluripotency network after the split of new world (NWM) and old world monkeys (OWMs). The divergence of PSC transcriptomes is also high among OWMs (Figures S5E-F). Compared to humans, we observe 2340, 375, 172 and 81 differentially expressed genes (DEGs) unique to *Callithrix*, *Gorilla*, *Chimpanzee* and *Bonobo* PSCs, respectively, whereas only 82 genes are differentially expressed in all the studied species (Figure S5E and Table S6). The number of unique DEGs is also directly proportional to the total number of DEGs, and the degree of transcriptome diversity agrees with the predicted evolutionary path as inferred from clustering fold change values of all observed DEGs (Figures 5C and S5E-G). As we expected given the evolutionary distance, the most contrasting transcriptional pattern is observed between the pluripotent cells of NWM and OWMs.

**Figure 5.**
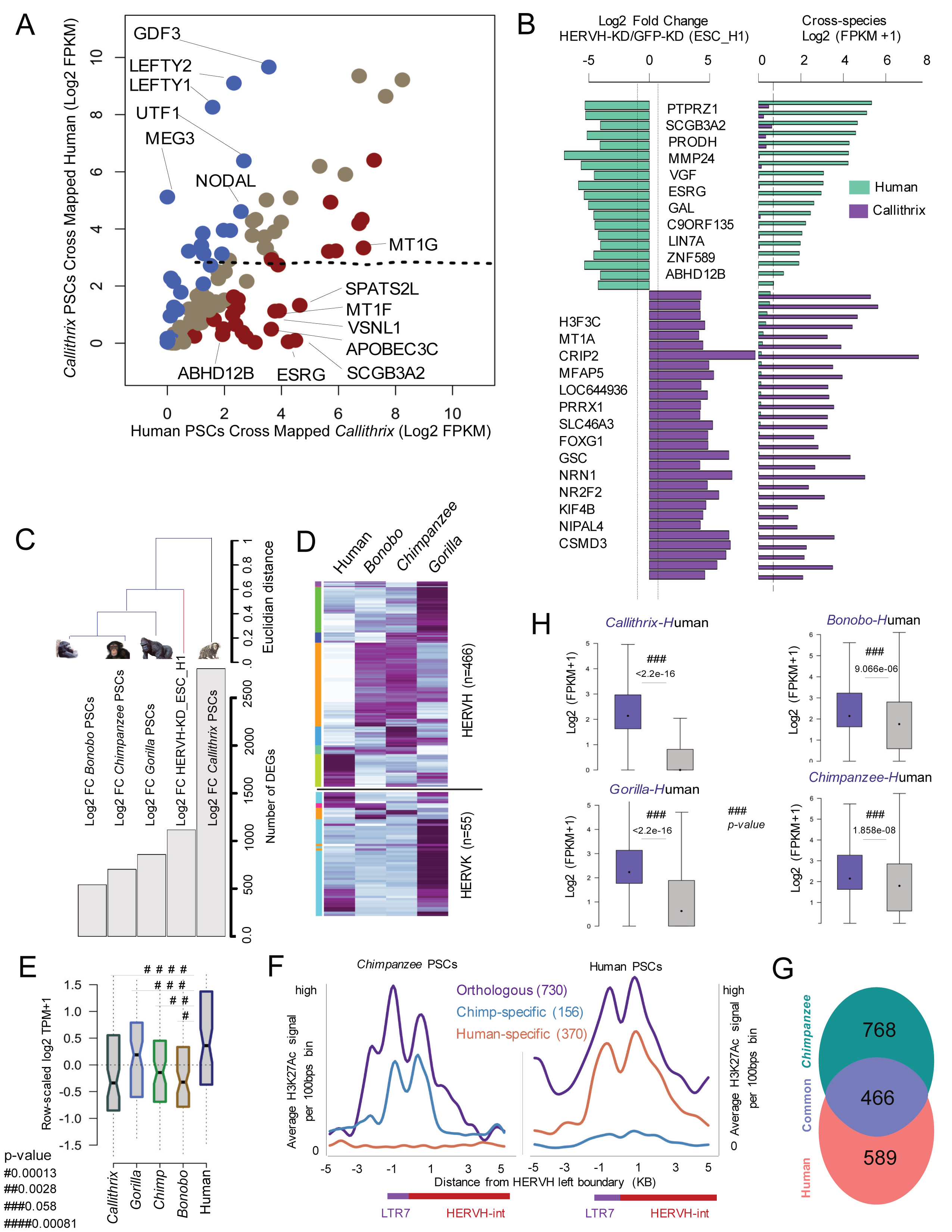
Species-specific gene expression in pluripotent cells during primate evolution is contributed by HERVH. A. Scatterplot shows differential expression of human pluripotent EPI genes (n=308, AUC cut off > 80%) in human and *Callithrix* pluripotent stem cells. Expression values are obtained as RPKM calculated on the human genome from reads mappable on to both genomes. Dots represent genes that have either lost (blue) or gained (dark-red) expression in human PSCs, respectively. Note: we also considered those genes that contained zero mappable reads in either of the analysed species (e.g. ESRG, SCGB3A2, APOBEC3C, etc). B. Combined barplots show the gain (n=19) and loss (n=29) of gene expression between human and *Callithrix* (no HERVH is present) pluripotent stem cells due to HERVH regulation. Left and right panels show fold-change |2| values in KD-HERVH vs KD-GFPcontrol in ESC_H1s and cross-species expression (RPKM), respectively. Green bars in the right panel are genes expressed significantly in human, but not in *Callithrix* (reads are mapped on to the human genome). Left panel shows the same set of genes downregulated when HERVH is depleted using by RNAi (HERVH-KD vs GFPcontrol-KD in ESC_H1s). The opposite scenario is shown in purple. (FPKM < 1 was considered as loss). C. Barplots combined with dendrograms display the comparison of genes in non-human primate and human pluripotent stem cells (PSCs) controlled by HERVH transcription. Barplots showing the number of significant differentially expressed genes (DEGs) (FDR<0.01 and fold change |2|) of gene lists obtained from *Callithrix* (n=1) vs human (n=4), *Gorilla* (n=2) vs human (n=4), *Chimpanzee* (n=4) vs human (n=4), *Bonobo* (n=4) vs human (n=4), HERVH-KD vs GFPcontrol-KD (n=2) in ESC_H1. In cases with two replicates, we selected only those genes, which were differentially regulated in both replicates in a similar fashion. Sorted according to HERVH-KD. The dendrogram representing hierarchical clustering is calculated using Euclidian distance method on the ranked correlation of most variable genes (n=~2800). D. The heatmap displays the dynamic expression of HERVH and HERVK (Log2 FPKM) in primate pluripotent stem cells. Z-scores are calculated on the expression obtained by mapping reads on the human genome from *Bonobo*, *Chimpanzee* and *Gorilla*. The numbers represent expressed genomic loci. Note the lower number of active genomic copies of HERVH exclusively in human compared to the other primates. E. Notched boxplots represent the distribution of relative expression levels of TR7/HERVH cis-target genes (STARR-seq, n=53) from cross-species mappable reads in various primate pluripotent stem cells (at Log2 scale). Hash (#) bars show the pairwise calculation of p-values between human and non-human primates. F. Active chromatin histone H3K27Ac marks around HERVH loci in human and chimp pluripotent stem cells. The Y axis is the normalized coverage of average ChIP-seq read counts per 100 bp bin. The X-axis shows the window of 10 KB consisting 100 bins each of 100 bps around full-length (FL) LTR7-HERVH-int-LTR7 transcription start sites (left boundary shown). Orthologous and species-specific loci were analysed independently and plotted together. Data show the mean H3K27Ac pattern over HERVH loci. Data from chimpanzee PSCs are shown in the left panel, human in the right panel. Note the orthologous and species-specific patterns. G. Venn diagram of the comparative analysis of orthologous and species-specific ChIP-seq peaks over HERVH loci that are active (occupied by H3K27Ac) in human vs chimp. H. Boxplots showing the pairwise distribution of global genomic expression of transposable elements (TrEs) between human (hg19 version) and non-human primate PSCs. We consider only those TrE loci that are mappable in both comparators and are expressed with an average Log2 FPKM > 1. Fidelity of comparison was checked using MA-plot for normalization of datasets (see Figure S5D). P-values were calculated by Wilcoxon test and adjusted using BH corrections.

How has ERVH expression changed over time? To examine this, we determine the expression of human HERVH loci by mapping reads from the comparator species against the human genome, and calculating the level of relative transcription at each locus. Among those genes whose expression has tuned down in human, we identified NR2F2, whose repression was reported to enhance PSC reprogramming (Hu et al., 2013) (Figure 5B). Curiously, PRODH is also among the HERVH-controlled gained genes, suggesting that PRODH is under a dual HERV-governed regulation e.g. LTR5/HERVK and LTR7B in brain (Suntsova et al., 2013) (Figure 5B), respectively. Differences between the NWM and OWM are also reflected in gene loss/gain expression events. Remarkably, the major gain (N=19) and loss (N=29) events between human and *Callithrix* are due to HERVH-governed gene expression (Figure 5B), underpinning the centrality of HERVH in turnover of the pluripotent cell transcriptomes in primates.

### The emergence of the HERVH-based regulatory network predates the human-gorilla common ancestor

When was the co-option of HERVH initiated? To address this, we employ the gene expression profile of the HERVH knockdown (HERVH-KD) as a surrogate of the ancestral – before HERVH – expression profile. Adding up all observed DEGs in the pairwise cross-species comparisons, including those identified in HERVH-KD, results in around 1100 genes (FDR < 0.05). Hierarchical clustering using ranked-correlation on their fold-change values reflects the evolution of the primate transcriptome, and pushes human HERVH-KD_ESC_H1 between *Gorilla* and *Callithrix* (Figures 5C and S5F-G), consistent with the domestication of HERVH predating the human-gorilla common ancestor.

How has ERVH expression changed during evolution? To examine this, we determine the relative transcription of each of the orthologous ERVH loci by mapping cross-species reads of the comparator species (gorilla and chimp) against the human genome. Compared to non-human (non-h) PSCs, we observe a heavy loss in the number of HERVH expressed loci in human PSCs, whereas only a few orthologous loci gained expression (Figures 5D and S5H-I). Curiously, a subset of orthologous ERVH copies appear to affect neighbour gene expression specifically in humans (Figures 5E), suggesting that the HERVH-mediated modulation of pluripotency, has occurred quite recently and some specifically in human.

To further consider the mechanisms of control and of turnover, we compare histone modification ChiP-seq (H3K27Ac) data on human and chimp orthologous and species-specific genome-wide active regions (Figure 5F). This strategy can detect the orthologous, as well as the species-specific, active ERVH loci. While, this analysis reveals peaks over active ERVH that are common to the two species (N~466), it identifies more peaks over specific loci that are unique to either chimp (N~700) or human (N~500), this being consistent with a recent ERVH-driven transcriptional remodelling process (Figure 5G). Curiously, upon comparing the orthologous transposable element (TrE) loci between primate species on normalized data (Figures S5C-D), we notice a marked reduction of overall TrE expression (including the Young, mutagenic elements) in humans, (Figure 5H) consistent with prior reports (Ward et al., 2018). This suggests that the establishment of the HERVH-driven modulation of pluripotency occurred in parallel (or included) the mitigation of overall TrE activities.

### Human naïve cultures are heterogeneous and are both evolutionarily and developmentally “confused”

Above we have characterised the various cell types in human pre-implantation embryos and shown that much of the circuitry is lineage-specific. In particular, the ERVH driven transcriptional network has significantly modulated pluripotency during primate evolution. How well do current *in vitro* pluripotent stem cell cultures match these features? We examine human naïve cell cultures (e.g. those with 3D morphology) that are either converted from primed cells (e.g. 2D morphology) or freshly established from the human blastocyst (Chan et al., 2013; Pastor et al., 2016; Takashima et al., 2014; Theunissen et al., 2016) and compare them and their potential primed counterparts. We expect the primed cells not to resemble the pluripotent *in vivo* cells and thus to be an ontogenetically-related negative control. The naïve cells should however resemble ICM/EPI.

Upon surveying expression of lineage-specific markers of the blastocyst (AUC cut-off > 85%) we observe that while the naïve cultures upregulate most of the ICM/EPI specific markers and downregulate PE-specific markers when compared to their primed counterparts, they also upregulate NCC markers (Figure 6A). To better profile them, we examine genes that are significantly expressed (log2 TPM > 1) in the cultured pluripotent lines (naïve and primed). Intersecting the resultant genes with our lineage specific markers (Figure 6A), we find that none of the cultured PSC lines express more than ~ 80% of the markers of any given lineage. These *in vitro* cultures appear to represent heterogeneous mixtures of cell types, some that express a range of pre-implantation lineage markers, some that are NCC-like (Figures 6A and S6A). Besides blastocyst, the current naive cultures also resemble 8-cell/morula (Theunissen et al., 2016), the phase at which youngest TrEs (e.g. LINE-1/SVA) are most active. In addition to transcriptomic similarity (Theunissen et al., 2016; Wang et al., 2016), this resemblance is also observed at the chromatin level. Indeed, both LTR5_Hs and SVA harbouring genomic loci exhibit a strong enrichment of ATAC-seq coverage at the 8-cell stage (Figure S6B), perhaps pointing to their potential regulatory role at this stage of development. By contrast, ATAC-seq signal intensity for LTR5_Hs/SVA genomic loci drops significantly in ICM, with a dynamic opposite to that of LTR7. Notably, in naive *in vitro* cultures the chromatin exhibits a similar status over SVA and hyperactive status over LTR5_Hs, but also at LTR7 loci (Figure S6B). Curiously, the hyperactive chromatin status is not necessarily reflected in transcript levels. In contrast to the upregulated Young TrEs, the number of transcriptionally active HERVH loci is low (N~50) in these naïve cultures (e.g. (Theunissen et al., 2016)), suggesting that most of the HERVH-driven transcripts, including those affecting human pluripotency (Wang et al., 2014), are not generated. This number is much lower when compared to HERVH loci at accessible chromatin regions in bulk ICM or PSCs (500 vs 250/300), potentially reflecting differences in HERVH-mediated pluripotency regulation in naive vs primed cells (Figures 6B and S6C-E).

**Figure 6.**
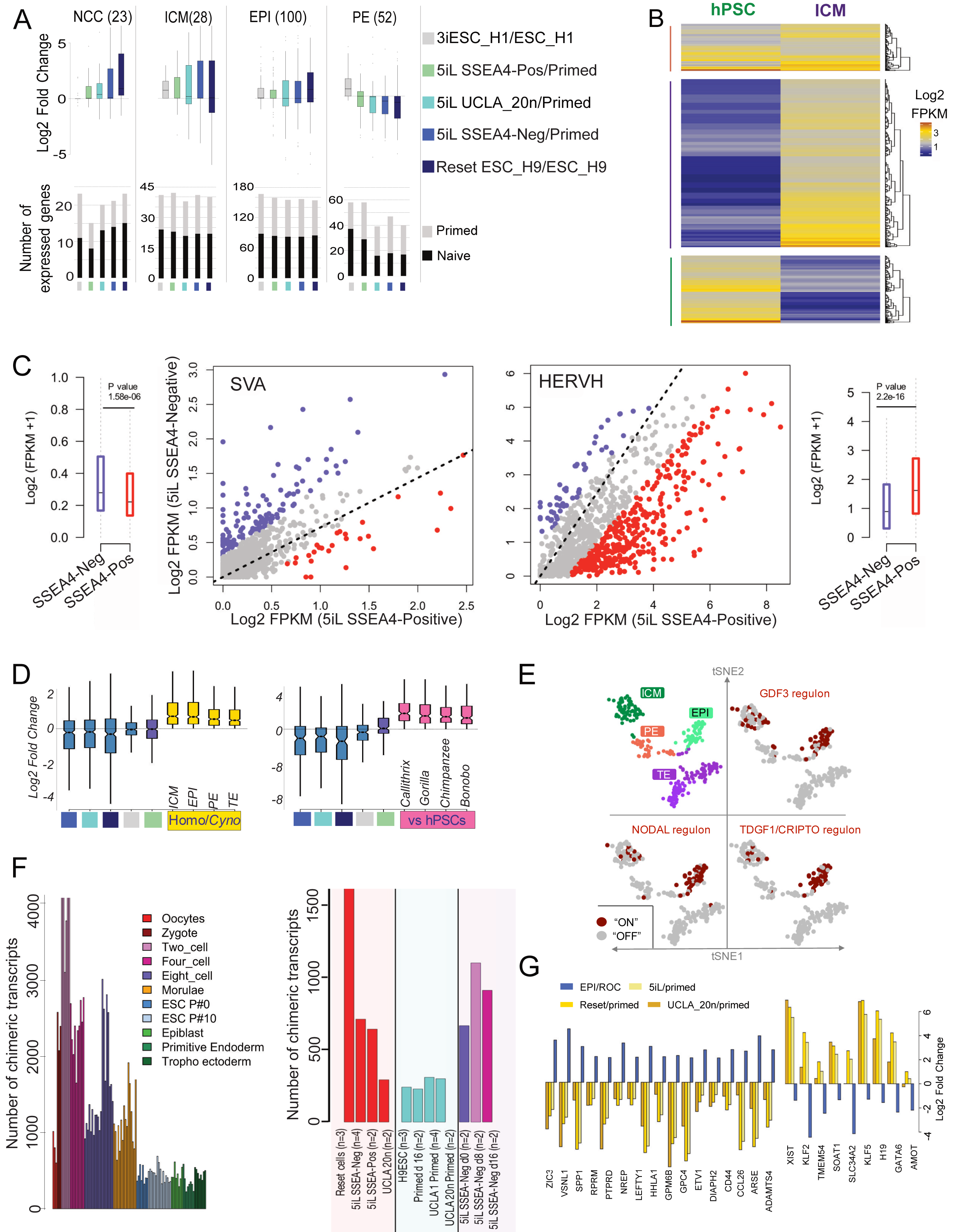
Lessons for *in vitro* models of early embryogenesis. A. Boxplots show the expression of human pre-implantation lineage markers in various naïve cell cultures compared to their respective primed counterparts (GFOLD calculated on Reset cells/ESC_H9s (Takashima et al., 2014; Theunissen and Jaenisch, 2014), 5iL_SSEA_Neg/UCLA1_primed, UCLA_20n/UCLA_20n_primed, and 5iL_SSEA_Pos/UCLA1_primed (Pastor et al., 2016). Note that the current naive cultures are heterogenous, as they express a variety of lineage markers. They upregulate ICM/EPI and downregulate PE markers, but express apoptotic markers of NCCs. The marker genes selected for the analysis flag distinct lineages (e.g. NCC, ICM, EPI, PE; N = number of marker genes used). The AUC cutoff values were chosen using the following criteria: the analysed gene is (i) a putative marker of any distinct lineage in human pre-implantation embryos (AUC cutoff > 0.85) after EGA; (ii) expressed in either naïve or primed cultures. Lower panels illustrate the number of lineage markers (AUC > 85%) expressed in the various *in vitro* cultures (naive and their primed counterparts) (Log2 TPM > 1). Note that the investigated naive cultures express less that 80% of any given lineage markers. B. Heatmap showing the comparison of ATAC-seq (Assay for Transposase Accessible Chromatin with high-throughput sequencing) enriched regions (Log2 FPKM > 1) over HERVH bins between bulk ICM and cultured hPSCs. Note that the while several HERVH loci exhibit a similar accessibility status in both hPSCs and the human ICM, the vast majority of the HERVH loci display differential coverage of ATAC-seq. C. Scatterplots showing differential expression of SVA (left) and HERVH (right) loci in naive cultures (5iL), sorted by a pluripotency marker (SSEA4). Boxplots next to the scatterplots represent the expression distribution of the respective TrE in SSEA4-sorted 5IL naive cells. Note the enriched HERVH expression in the cells sorted for a pluripotency marker. D. Current naive cultures under-represent human-specific features. Notched boxplots showing the distribution of differential gene expression (Log2 fold change) in various human naïve cell cultures with their respective primed counterparts (steel blue boxes). The genes chosen for the comparative analysis (N=246) presented on the left panel are upregulated (Log2 fold change > 1) in all lineages (ICM, EPI, PE and TE) of the human (*Homo*) blastocysts compared to their counter lineages in macaque (*Cyno*) blastocysts (gold boxes). Note that these genes, representing the “human specific features” of blastocyst evolution since macaque (gold boxes) are downregulated in the various naive cultures (steel blue boxes). The genes chosen for the comparative analysis (N=197) presented on the right panel are upregulated (Log2 fold change > 1) in human pluripotent stem cells (hPSCs) vs all analysed non-hPSCs (*Callithrix*, *Gorilla*, *Chimpanzee* and *Bonobo*) (violet boxes) (Right panel). Note that these genes, reflecting “human specific pluripotent stem cell features” compared to non-human primates (violet boxes), are downregulated in the studied naive cultures (steel blue boxes). E. tSNE clusters on the expression matrices of the human blastocyst coloured with respect to lineages (ICM, EPI, PE and TE) (upper left). Top gene regulatory networks (regulons) of the EPI state predicted by SCENIC (e.g. driven by NODAL, GDF3, TDGF1/CRIPTO master regulators). Feature plots, based on tSNE plot show the binary activity matrices after applying SCENIC. Red and grey dots represent single cells where the respective regulons are active or inactive (see also Figure S4B). Note: Self-renewal regulons are co-expressed in most of the EPI cells, but not in PE or TE clusters. F. Barplot showing the number of chimeric transcripts detected in single cell transcriptomes of distinct human pre-implantation stages (Yan et al., 2013) (left panel). Note that the generation of chimeric transcripts reduced after morula. Bar plot showing the number of chimeric transcripts detected in various naïve cultures, in their respective primed controls and in their converted counterparts (N = number of samples analysed) (right panel). G. EPI checklist. Log2-fold change values of genes either specifically expressed in pre-EPI/EPI vs rest of the cells (ROC) in the human embryos, but showing the opposite pattern when naïve cells are compared with their primed counterparts.

The current naïve cultures exhibit a non-stereotypical idiosyncratic expression profile, and are in this sense developmentally “confused”. The overexpression of potentially mutagenic TrEs (expression which may under normal circumstances lead to apoptosis and a sideways move into the NCC category) might be of safety concern for therapeutic application of these *in vitro* cultures or for their long-term stability. Enriching naive cultures for pluripotency (e.g. by sorting by SSEA4) improves quality (Pastor et al., 2016). Importantly, besides positive decreasing heterogeneity of the culture, SSEA4^positive^ cells show high expression of LTR7/HERVH and mitigated SVA expression (Figures 6C).

Are the transcripts that mark the human-specific features of pluripotent cells *in vivo* well represented in naïve cultures? To address this, we also compared fold-change expression of naïve vs primed cells with the fold-change expression of pairwise human and macaque blastocyst stages. Our analysis reveals that the human pluripotent blastocyst features are, surprisingly, significantly downregulated in naïve cultures (Figure 6D). Similarly, transcripts of those genes that mark the human specific vs primate features of pluripotent stem cells are underrepresented in naïve cultures (Figures 6D and S6F). In this regard, the naïve cells are also phylogenetically “confused”.

In addition to being developmentally and phylogenetically confused, in some regards they also don’t convert to primed cells in the expected manner (Pastor et al., 2016). Chimeric transcripts (transcripts deriving from two physically independent genomic loci) are abundantly generated until and including the morula stage, but their generation gradually decays to an approximately steady-state frequency after (Figure 6F). Unexpectedly then, chimeric transcripts are still expressed upon converting naïve cells to primed state (Figure 6F). This might go some way to explaining why these cells are less capable of proper conversion (Pastor et al., 2016).

### Defining the optimal *in vitro* cell type

Understanding early human embryogenesis could also help to define a cell type that would be a good candidate for a laboratory model pluripotent cell population. One of the criteria is pluripotency, thus ruling out PE this being a committed lineage. Given the expression of Young mutagenic TrEs in morula, we focus interest on ICM and EPI. We analysed the gene expression pattern of pluripotency-specific markers (e.g. NANOG, KLF4, POUF5F1/OCT4, etc.) in both. We find no differential pluripotency gene expression between EPI and ICM (p-value insignificant) (Figure S6G). Thus, EPI and ICM are both pluripotency candidates, arguing against EPI being the only pluripotent cell population in the pre-implantation embryos (Brons et al., 2007).

Optimally, in addition to pluripotency, the cultured cells would require self-renewal ability to maintain a relatively homogeneous and genetically stable cell population. Notably, among the genes that could be regarded as top markers defining EPI against ICM, (e.g. NODAL and GDF3; rank 1 and 2 in our analysis, Figures 1A, S1F and S6G) are those implicated in triggering the self-renewal cascade (Niakan and Eggan, 2013). In addition to NODAL and GDF3, our SCENIC analysis identifies further genes associated with self-renewal (e.g. LEFTY1/2, TDGF1, TDGF1P3) in the reconstituted top regulon of EPI (Figure S4B-D). Expression profiling of these self-renewal genes indicates that the self-renewal potential is a property of EPI, but not of ICM (Figure S6G). These results define EPI as a pluripotent cell population with self-renewing hallmarks and hence a most appropriate candidate for *in vitro* work.

If EPI is possibly the best cell type to mimic *in vitro*, what genes should the optimal cell type express or not express? To this end, we propose a checklist that could be used to guide *in vitro* studies. We presume that the key properties of the pluripotent EPI, including self-renewal and homogeneity, make this the best candidate to mimic *in vitro*. The checklist hence includes top genes that appear to have a higher expression status in human EPI when compared against the other clustered cell types (Figure 6G). We also provide a checklist to exclude genes that are downregulated in EPI, and do not feature in any developmental stage in human (Figure 6G). The expression of these genes could induce various aberrant processes that could compromise pluripotency. The checklist of included genes features ESRG, LEFTY1, HHLA1, while excluded genes include H19 and KLF2/5 (Figure 6G). Notably, ESRG and HHLA1 genes carry full-length HERVH sequences, suggesting that human pluripotent cultures should reflect the species-specific features properly, including the expression of HERVH-remodelled genes that are associated with pluripotency regulation in humans.

## Discussion

To discern the uniqueness (or lack thereof) of human early development, we aimed to decipher both the structure and the evolution of the blastocyst, with a special focus on pluripotency regulation in primates. Despite prior assumptions, we find that the trajectories and the cell types in the primate blastocyst are fundamentally conserved. Adding to the standard hourglass model of development (Kalinka et al., 2010) that notices rapid transcriptome evolution of early cell types, we find that EPI is highly divergent between humans and non-human primates, but gene expression in other cell types (ICM, PE) is relatively conserved. The major HERVH-driven transcriptional remodelling of EPI has occurred quite recently, following the split of the *Gorilla*-human common ancestor, accompanied by a heavy loss of expressed HERVH orthologous loci and gain of species-specific active loci. Thus, pluripotency of the human form appears to be specific to us humans.

Concurrent with unmasking ICM, we identified a previously unrecognised cell population during blastocyst formation, NCCs, that are exposed to apoptosis. Unless recognised, these NCCs interfere with the proper definition of ICM and explain why prior putative ICM populations expressed apoptotic genes (e.g. in (Stirparo et al., 2018)). In turn the definition of the derivative EPI and PE markers is confused. Apoptosis is thought to play an active role throughout the developmental process. Apoptotic cells are first detectable immediately following embryonic genome activation (EGA), but their timing varies among different mammalian species (Braude et al., 1988). In the mouse, the major activation event occurs during the 2-cell stage (Flach et al., 1982), whereas in humans it occurs later, between the 4-and 8-cell stages (Braude et al., 1988). EGA is accompanied with a global epigenetic change that also de-represses transposable elements (TrEs) (Rowe and Trono, 2011) and this, we suggest, may explain the appearance of NCCs. In human, the expression of phylogenetically young elements peaks in morula (Romer et al., 2017; Theunissen et al., 2016). While certain Young TrE (e.g. SVA, LTR5_HS) copies might have a regulatory function following EGA, the genomic loci of Young TrEs are typically inactivated during the morula-blastocyst transition. In morula, where asymmetrical cell division already generates diversified epigenomes, upregulated Young TrEs are likely to further contribute to the observed high heterogeneity of cells, giving rise to both committed (progenitor) and non-committed cells (NCCs). It is parsimonious to suppose that apoptosis is a means to enforce a selective filter against damaged cells that emerge after EGA and fail to properly express lineage markers. A large population of morula-derived NCCs are not observed in mice, perhaps due to the earlier timing of EGA, while they might exist in primates, but went undetected (e.g. owing to selective cell extraction (Nakamura et al., 2016)).

According to our quality control model, pre-implantation development is not absolutely directional, but includes a selection process. The boost of TrE activity might determine the fate of the embryo as a whole, whether it proceeds to the blastocyst stage or be selected out in a process reminiscent of attrition in fetal oocytes (Malki et al., 2014), involving programmed cell death. If the embryo were to be viable, cells that cannot control mutagenic TrEs and fail to commit would be targeted to apoptosis, whereas cells capable of mitigating transposition activities and express lineage markers would develop further. Nevertheless, it is likely that some TrE activity is tolerated, resulting in TrE integration in the trophoblast (see also Muñoz-Lopez. et al., 2018), and even in heritable TrE insertions accumulated through transposition at this stage of human development (van den Hurk et al., 2007). An alternative possibility is that NCCs express mutagenic TrEs because they have no developmental fate. In this model, the upregulation of young TrEs could be a mechanism to lead to the destruction of cells that, for whatever reason, have failed to commit or that are surplus to requirements. Consistent with both models, NCCs are not detectable after E5, thus excluded from the developmental process.

### Do pluripotency and host-defence coevolve?

The connection between pluripotency and TrEs can also go some way to explaining why the evolution of pluripotency in primates is associated with the innate immunity and defence response. The connection between the evolution of pluripotency and self-defence has been suggested before (Grow et al., 2015; Wang et al., 2014) to explain why the human defence response was capable of dealing with the multiple waves of viral invasion during primate evolution with successful attenuation of Young TrEs (Friedli and Trono, 2015). From the arsenal of complementary processes regulating TrE/viral activities we have detected various APOBECs (Knisbacher et al., 2016) and IFITM1, specifically expressed in ICM/EPI (rather than NCCs). Intriguingly, in contrast to NCCs that are marked by Young TrEs, progenitor cells that passed quality control and continue to participate in the developmental program characteristically express relatively old, dominantly full-length ERVs, primarily HERVH. The phenotype of Young TrE activation can be reproduced in HERVH-knockdown conditions in pluripotent stem cells (Lu et al., 2014), suggesting that relatively old TrEs might be involved in Young TrE suppression. This hypothesis is strengthened by our finding of possible HERVH-mediated regulation of APOBEC3C/D/G. If so, this would suggest an interesting case history in which one TrE is retaining owing to its ability to suppress others.

HERVH appears to selectively mark the genetically stable, self-renewing, pluripotent cell population during blastocyst formation. Why, evolutionarily speaking, might this be? HERVH seems rich in binding sites for pluripotency transcription factors not just through the LTR but also across the full gene body. One can speculate that successful endogenization of the ancestral retrovirus H was dependent on its ability to use TFs that are present in the target cell type of the endogenization (i.e. not somatic cells, but pre-germline cells). The net consequence of a successful process is the distribution of multiple endogenized retrovirus copies by transposition with an inherent ability to drive novel gene expression in their near vicinity in key pluripotency cells. Some of these transcripts may well be transcriptional noise, but some seem to have become engrained into the pluripotency circuitry. Inclusion into such a circuitry is consistent with a model in which TrEs are under selection to control cell fate to promote their own “ends”, and thus to maintain pluripotency or to steer cell fate towards germline, within which transposition events have an evolutionary future (Izsvak et al., 2016).

### Lessons for *in vitro* models of early embryogenesis

In the last five years, numerous attempts have been reported to derive human naive cell cultures. The quality of these cells has been extensively discussed, e.g. (Theunissen et al., 2016, Pastor et al., 2016; Takashima et al., 2014). Researchers aim at establishing pluripotent stem cell cultures, mimicking the pluripotent blastocyst as closely as possible, and at finding a suitable non-human primate host for human pluripotent stem cells. *Cynomolgus* is assumed to have both a comparable pluripotency regulation and to serve as a potential *in vivo* model of human biology (Dodsworth et al., 2015).

Our analysis holds lessons for establishing self-renewing, pluripotent stem cell cultures *in vitro*. The human pluripotent EPI forms a relatively homogenous cell cluster, has self-renewal hallmarks and attenuated Young TrE activity, potentially making a good choice for *in vitro* culturing. While, the studied naïve cultures have improved presentation of ICM/EPI markers and underrepresent PE markers, they are heterogeneous, each consisting of a mixture of cells of various identities, and appear to be both evolutionarily and developmentally “confused”. Beside EPI-like cells, naïve cultures contain large numbers of NCC-like cells, as well as cells that display similarity to various embryonic cell lineages, and to pluripotent cells of NHPs instead of human ones. Whether the human-specific features of EPI could explain difficulties faced by cross-species chimeric studies is yet to be clarified. Furthermore, as the desired end-point of the naïve-like cultures is as a model for early human development, or for transformation into any number of alternative cell types for therapeutic application, it is a matter of substance to discern whether potentially damaging transposition is happening. Filtering out NCCs, preserving human- and lineage-specific features should help to improve derivation and maintenance of human naïve stem cell cultures with improved homogeneity, pluripotency and genome stability. While several strategies have been suggested to establish naive-like cultures, employing a LTR7/HERVH reported-based approach (Wang et al., 2016; Wang et al., 2014) targets to identify pluripotent EPI-like cells, the only pluripotent self-renewing cell type in the pre-implantation embryo, could have multiple advantages. We suggest that inhibition of BMP2 or PDGFRA, co-expressed on the trajectory of ICM to PE, might lead to self-renewing EPI-like cells *in vitro*.

Commonly we employ mouse as a model for early human development. In this context it is important to note that self-renewal appears to be a hallmark of human EPI, as murine EPI shows no similar self-renewal hallmarks. In retrospect this difference may be expected.

While the human EPI is formed at E5 and maintained as EPI between E5 and E7 as a population of cells with highly similar transcriptomes (i.e. they can self-renew), in mouse, by contrast, EPI is short lived with the embryo being implanted immediately after EPI’s generation. Other human cell types of the blastocyst (e.g. PE and TE) are more similar to mouse EPI in that they differentiate from early to late stage during the time course over which EPI is maintained. Human EPI thus appears unique as a cell type, both in comparison to mouse EPI and other human cells, requiring relative long-lived homogeneity. Involvement of genes like HERVH-remodelled ESRG, SCGB3A2, ABHD12B in the EPI-specific module reinforce the human-specificity of EPI self-renewal and underscore the importance of understanding the phylogenetically distinct nature of early human embryogenesis if we are to optimize *in vitro* cultures.

## Methods

### Bulk RNAseq

#### Data generation

Total RNA was extracted from *Callithrix jacchus* (Muller et al., 2009) and Gorilla PSCs (Wunderlich et al., 2014) using Trizol RNA Mini Prep kit (Zymo research) following the manufacturer’s instructions. Following RNA extraction, DNase treatment was applied using TURBO DNA-free Kit (Ambion). The RNAseq library preparation followed the Illumina TruSeq Stranded mRNA Sample Preparation Kit protocol on Illumina HiSeq machine with paired-end 101 cycles.

#### Data selection

Seventeen data sets were used for the analyses. In order to dissect the human pre-implantation lineages, we re-analyzed single cell RNAseq datasets from *Homo sapiens* (GSE36552 and E-MTAB-3929) and *Cynomolgus* pre-implantation embryogenesis (GSE74767). We used only predefined ICM, EPI, PE and TE samples from *Cynomolgus*, in order to perform comparative analysis with human counterparts. We also analysed mouse blastocyst samples (GSE45719 and GSE57249) to check the self-renewal networks in EPI (data not shown). We used ATAC-seq and RNAseq datasets from human 8-cell, bulk ICM (ICM/NCC mix), naïve and hESCs (GSE101571) and ChIP-STARR-seq, ChIP-seq of NANOG, POU5F1, H3K27Ac and H3K4Me1 from cultured naïve and primed hPSCs (GSE99631, GSE54471 and GSE35583) to decipher the HERVH-target genes in human pluripotency. To perform the comparative analysis of primate pluripotent states, we utilized human, Chimpanzee, Bonobo (GSE47626), Gorilla and Callithrix (this study). To compare the chromatin status at HERVH loci in primate pluripotency by histone modifications, we used H2K27Ac and H3K27Me3 marks in human and chimp PSCs (GSE69919). We did not find significant enrichment of H3K27Me3 over HERVH in either of species, so do not present it. To analyse the pluripotent states cultured in different conditions, we obtained the RNAseq datasets of 3iL (E-MTAB-2031), 5iL (SSEA-Neg, SSEA-Pos), UCLA20n (GSE76970), Reset, HNES (E-MTAB-2857, E-MTAB-4461) naïve and primed cells, and compared them with HERVH-KD datasets in ESC-H1 (GSE38993) in pairwise manner.

#### Data processing

RNAseq reads with MAP quality score < 30 were removed. We checked the quality of sequencing reads using FASTQC (https://www.bioinformatics.babraham.ac.uk/projects/fastqc/). We also truncated 2-5 nt from the end of sequencing reads, since their average quality score was lower than that of the rest of the sequence. This resulted in at least 70 million reads per sample of *Gorilla* and *Callithrix*. Next, we mapped the reads over the reference genome (Human hg19/GRCh37) and transcriptome model (hg19.refseq.gtf) using STAR, downloaded from USCS tables (http://hgdownload.cse.ucsc.edu/goldenPath/hg19/bigZips/). First, STAR genome/transcriptome indices were constructed given their respective RefSeq gtf annotations. The reads were mapped to their respective reference genomes i.e. human (hg19), chimp (PanTro4), gorilla (gorGor4) and marmoset (calJac3) using STAR with our defined settings i.e –*alignIntronMin 20 –alignIntronMax 1000000 –chimSegmentMin 15 – chimJunctionOverhangMin 15 –outFilterMultimapNmax 20.* As per STAR default, we permitted at most two mismatches per 100 bp. Mapping quality of sequencing reads were calculated using RNA-SeQC (*https://software.broadinstitute.org/cancer/cga/rna-seqc*). The read counts were calculated with *featureCounts* from the *subread* package (http://subread.sourceforge.net/) at gene level with RefSeq annotations of genes and repeat elements. Gene expression levels were calculated as transcripts per million (TPM) from counts over the entire gene (defined as any transcript located between transcription start (TSS) and end sites (TES)). FPKM was calculated using *bamutils* (http://ngsutils.org/modules/bamutils/count/) for genes and repeat elements. Single cell RNAseq datasets were also re-mapped and expression was calculated for genes (Log2 TPM) and repeat elements as counts per million (CPM). In order to calculate differential expression at the gene level (bulk RNAseq), we used the GFOLD (https://zhanglab.tongji.edu.cn/softwares/GFOLD/index.html) algorithm, which calculates the normalization constant and variance to extract fold changes from unreplicated or unequally replicated RNAseq data. Differential expression of repeat elements was calculated at relative expression level which was further compared in pairwise manner. P-values were adjusted for False Discovery Rate (FDR) using BH corrections.

### Cross-species analyses

Genes that are differentially expressed (DEGs) between species were obtained by cross-species mapping of RNAseq reads. Reads mappable on both comparators were further mapped on the human genome (hg19) using STAR. Cross-species read counts, FPKM and effective fold-change were calculated using the tools mentioned above. To determine the global expression level of human genes and repeat elements in NHPs, we mapped human and non-human iPSC RNAseq reads against the human reference genome and gene models. Further, we used mappable reads over orthologous gene and repeat elements to perform pairwise comparative analysis.

### Single cell RNAseq data processing

Activity of genes in every sample was calculated at TPM expression levels. We considered samples expressing more than 5000 genes with expression level exceeding the defined threshold (Log2 TPM > 1). We considered genes expressing in at least 1% of total samples for the analysis. This resulted in 1285 single cells of human E3-E7 samples, with 15501 expressed genes. We used Seurat 1.2.1 for clustering E3-E7 and E5 cells (this version was the latest during the preparation of the manuscript) whereas the rest of analysis was carried out using Seurat_2.2.1 (http://satijalab.org/seurat/) packages from R to robustly normalize the datasets at logarithmic scale using “*scale.factor = 10000*”. After normalization, we calculated scaled expression (z-scores for each gene) for downstream dimension reduction. The cells were separated by subjecting the MVGs ({log(Variance) & log(Average Expression)} > 2) to the dimension reduction methods of principal component analysis (PCA).

### Principal component analysis (PCA)

As previously described (Macosko et al., 2015), we ran PCA using the ‘*prcomp*’ function in R, then utilized a modified randomization approach (‘*jack straw*’), a built-in function in the “Seurat” package, to identify ‘statistically significant’ principal components in the dataset. We used the genes contributing to the top 9 significant principle components (PCs) for E3-E7 stages, 5 significant PCs for E3-E5 stages, 3 significant PCs for E3-E4 and the first two significant PCs for E5 stage as input to visualize in 2D with tSNE.

### t-stochastic neighbour embedding (tSNE)

Using the above significant PCs as an input, we applied *tSNE*, a machine learning algorithm to cluster the cells in two dimensions. To define cell population clusters, we employed the *FindClusters* function of “Seurat” using “PCA” as a reduction method. To resolve the clusters on tSNE, the density parameter {G.use} was set between 6 to 10, and the parameter providing the fewest clusters was chosen for visualization. This approach identified 10 clusters from E3-E7, 5 clusters from E3-E5, 3 clusters from E3-E4 and E5 stages. The specific markers for each cluster identified by “Seurat” were determined by the “FindAllMarkers” function, using “roc” as a test for significance. A gene matching the following criteria was considered as a marker for a given cluster: (i) the gene is overexpressed in that particular cluster (average fold difference > 2 compared to the rest of the clusters), (ii) the gene is also expressed (Log2TPM > 2) in at least 70% of the cells in that particular cluster and (iii) the Area Under Curve (AUC) value is greater than 80%.

Feature plots, violin plots and heatmaps were constructed using default functions, excepting the colour scale that was set manually. The annotated ICM-EPI-PE cells (Figure 1A and 1E) were re-clustered using the methodologies described above and visualized on the tSNE coordinates.

### Trajectory and diffusion analysis

Trajectory analysis of the differentiation process from progenitors to committed cells was performed using the Monocle2 package (Qiu et al., 2017), which generates a pseudotime plot that graphically illustrates the branched and linear differentiation processes. For pseudo-temporal analysis of human and macaque data, we first imported the processed Seurat object into the Monocle2 workspace using “importCDS” function. The datasets were processed further using the series of default functions with negative binomial expression family parameters. The final dimensionality was reduced to two components. The dimensionality of the data was reduced by constructing a parsimonious tree using “DDRTree”. We employed the “differentialGeneTest” function to find the top DEGs (q-value < 1e^−8^), these being fed as an input for unsupervised ordering of the combined set of cells. Note: the q-value threshold was set manually (10^−8^ − 10^−15^) to find the root, branching point and leaves on the trajectory graphs. This approach identified the genes (100 to 1000) that were significantly differentially expressed. We then used the expression data of these genes to construct a diffusion map for the respective cell populations (DiffusionMap function in the Destiny package (Angerer et al., 2016)). We calculated the diffusion pseudo-time (DPT function in the Destiny package). Finally, the cells on the diffusion maps and trajectories were annotated on the basis of their identity, previously determined from the Seurat analysis. A similar strategy was also used on data from E5, E3-E5 and ICM-EPI-PE cells. Genes were plotted on a branched heatmap obtained by setting a threshold of differential expression to qval < 1e-8.

### Analysis of repetitive elements

To estimate expression levels for repetitive elements, we used two strategies. The long reads in Yan et al.’s data (Yan et al., 2013) allowed us to calculate CPM or RPKM for individual TrE loci. In contrast, data from (Petropoulos et al., 2016) was suitable only to detect the average expression of any given TrE family as it was unable to unambiguous map to any given locus exclusively. In this instance, we considered multimapping reads only if they were mapping exclusively within a TrE family. One alignment per read was employed to calculate counts per million (counts normalized per million of total reads mappable on the human genome). The expression level of repeat families was calculated as Log2 (CPM+1) prior to comparison. SVA-D elements from NCC and HERVH-int elements from EPI were the most abundant in the respective cell types. Given this, we removed the very few single cells that showed extremely low or non-detectible expression of these in the relevant cell type (Log (CPM+1) < 0.1). Note that datasets from different layouts (single vs bulk RNAseq) were never merged into one data frame to perform TrEs comparative analysis, not least because we were unable to normalise these datasets.

### ATAC-seq data analysis

ATAC-seq raw datasets in sra format were downloaded and converted to fastq format using *sratools* function *fastq-dump-split-3*. Fastq reads were mapped against the hg19 reference genome with the bowtie2 parameters: –very-sensitive-local. All unmapped reads, reads with MAPQ < 10 and PCR duplicates were removed using Picard and *samtools*. All the ATAC-seq peaks were called by MACS2 with the parameters –nomodel -q 0.01 -B. Blacklisted regions were excluded from called peaks (https://www.encodeproject.org/annotations/ENCSR636HFF/). To generate a set of unique peaks, we merged ATAC-seq peaks using the *mergeBed* function from bedtools, where the distance between peaks was less than 50 base pairs. We then intersected these peak sets with repeat elements from hg19 repeat-masked coordinates using bedtools *intersectBed* with 50% overlap. This provided 14203 peaks for Reset cells, 17439 for 5iL Naïve cells, 12028 and 11724 for Eight cell replicates, 6805 and 7505 for hESC replicates, 7270 and 3084 peaks for bulk ICM cells. If these peaks fell in a window of +/- 10KB around the TSS of protein-coding genes then they were assigned as a probable enhancer. SVA, LTR5 and LTR7 were found to be the Repeat families harbouring the highest number of peaks in the analysed samples. In order to calculate the enrichment over the given repeat elements, we first extended 2KB upstream and 8KB downstream coordinates from the left boundary (TSS) of respective elements in a strand-specific manner. These 10KB windows were further divided into 100 bps bins and tags (bedGraph output from MACS2) were counted in each bin. Tag counts in each bin were normalized by the total number of tags per million in given samples and presented as Counts Per Million (CPM) per 100 bps. CPM values were averaged for each bin between replicates prior to plotting the figures. Note: Pearson’s correlation coefficient between replicates across the bins was found to be r > 0.85.

### SCENIC data analysis

This computational strategy uses the GENIE3 algorithm (Aibar et al., 2017). Inferring Regulatory Networks from Expression Data Using Tree-Based Methods) to construct modules, these being gene sets that are co-expressed with a master regulator, using the random forest regression. SCENIC provides several thresholds to build valid modules. We only selected the top 50 targets with the highest weight for each TF (links with weight > 0.01) to avoid an excess of arbitrary thresholds. Only the gene sets (co-expression modules) with at least 20 genes were kept for the AUCell scores (with *aucMaxRank* = 10%). We used either the AUC scores directly for a heatmap, or a binary matrix using a cutoff (determined automatically) of the AUC score.

### Orthologous and species-specific TrE analysis

The hg19 RepeatMasker track (http://www.repeatmasker.org) was converted to panTro4 and gorGor3 genomic coordinates using liftOver reciprocal chain files (http://hgdownload.cse.ucsc.edu/goldenPath/hg19/vsPanTro4/reciprocalBest/ and http://hgdownload.cse.ucsc.edu/goldenPath/hg19/vsgorGor3/reciprocalBest/), requiring a minimum 70% sequence match. These genomic coordinates were intersected with the hg19 *RepeatMasker* track, requiring that 50% of the base pairs overlap with the similarly annotated sequence in the comparator species. These coordinates were further lifted back to hg19, and considered as orthologous TrEs. Species-specific TrEs for each species were identified after excluding the orthologous TrEs (e.g. the TrE name is not present in the *RepeatMasker* track of the other species).

### Cross-species ChIP-seq data analysis

ChIP-seq reads of H3K27Ac, H3K27Me3 and total input from human and *Chimpanzee* PSCs were mapped to their respective genomes and across the genome (reciprocally to hg19 and panTro4) using *Bowtie 2* (Langmead and Salzberg, 2012) with the –*very-sensitive* option and mapping quality filter of MAQ > 10. Peaks were called using MACS2 with the –*broad -g hs –broad-cutoff 0.1* options. A threshold of *qvalue* < 0.01 was used in further analysis. Peak co-ordinates from both species were then converted to their counter species genome using liftOver and the best reciprocal chain (http://hgdownload.cse.ucsc.edu/goldenPath/hg19/vsPanTro4/reciprocalBest/) between hg19 and panTro4 requiring a minimum 70% sequence match. Peak regions that could be reciprocally and uniquely mapped between the two genomes were considered as orthologous peak regions. Human-chimp orthologous HERVH regions were obtained using the LiftOver tool to cross-species mapping over the “best reciprocal chain alignments” (http://hgdownload.cse.ucsc.edu/goldenPath/hg19/vsPanTro4/reciprocalBest/) between hg19 and panTro4, requiring 70% sequence match. ChIP-seq reads were mapped on both genomes. Those unmapped sequences which were annotated as HERVH by *Repeatmasker* in either of the species were considered to be Human/Chimpanzee-specific.

### Identifying HERVH target genes

We determined HERVH-derived functional enhancers in human pluripotent cultured stem cells (hPSCs) by data mining the merged analysis of ChIP-seq, plasmid DNA-seq and ChIP-STARR RNA-seq (Barakat et al., 2018)(GSE99631). Using these genome-wide analyses, we first collected HERVH loci with RPPM >144 reads per plasmid million (RPPM). This strategy identified a list of 543 distinct HERVH loci as functional enhancers in hPSCs (STARR-seq was performed on both naive and primed cell types). In addition, we aimed at determining open chromatin regions around HERVH loci from bulk ICM cells (Note: this was a mixture of ICM and NCCs) (Wu et al., 2018). First, the signal file (wig format) of ATAC-seq in ICM and hESCs was downloaded and RPKM values were calculated per 100⃠bp-window. As previously (Wu et al., 2018), windows carrying RPKM > 2 were considered as open chromatin regions. Next, we intersected these regions with the 100 bp bins of distinct full-length HERVH loci, resulting in a list of HERVH loci at accessible chromatin regions (in the case of multiple bins overlapping within a HERVH loci, the bin with the highest RPKM was considered). This resulted in 569 distinct HERVH loci serving as either enhancer or promoter in bulk ICM. We hypothesized that the HERVH regions commonly identified in hPSCs (both in *in vitro* and *in vivo*) could be considered as “universal” pluripotent HERVH-derived enhancers. We found 282 HERVH accessible chromatin regions (ATAC-seq) overlapping with STARR-seq peaks identified in cultured hPSCs. Among these 282 loci, we identified 72 loci located in the vicinity of a gene (defined as no more than 10 kb distant from the TSS of genes). We considered a neighbour gene as a HERVH-target when no additional ATAC-seq peaks could be detected between the HERVH and the TSS. Using this strategy, we catalogued 43 HERVH-target genes in the human ICM.

### Enhancer strength of TrE-derived sequences

We leveraged the advantage of available genome-wide functional enhancer (STARR-seq) datasets in naïve and primed cells from GSE35583. We first intersected hg19 repeat masked coordinates with enhancer coordinates using bedtools intersectBed with 50% overlap. To generate a set of unique enhancers per repeat locus, we next merged enhancer peaks using mergeBed function for which the distance between peaks was less than 50 base pairs. Enhancer strength of a repeat family was calculated as n×1000000/NT, where n = number of TrE loci within its family that overlap with ChIP-STARR-seq regions; N = Total number of TrE loci within a family in human genome; T = Total number of enhancer regions obtained in human genome. Note: High and low enhancer regions were catalogued previously (Barakat et al., 2018).

### *Homo-Cynomolgus* cross-species comparative analysis

For this cross-species analysis, we selected 228 cells from human pre-implantation blastocysts (Petropoulos et al., 2016) (ICM, EPI, PE and TE) and 170 cells from *Cynomolgus* (Nakamura et al., 2016) (ICM, EPI, Hypoblast (PE) and TE). For generating a cross-platform single-cell RNAseq dataset, counts were merged by gene name, and log2 TPM+1s were calculated. We redefined ICM, EPI, PE and TE cells using only those genes that were annotated in Refseq gene tracks of both human and *Cynomolgus* (similar in approach to Nakamura et al., 2016). This approach resulted in 16222 individual genes that were merged into a single pool. We restricted the analysis to 11053 orthologous genes that were expressed (Log2 TPM > 1) in at least 5 cells out of the ~400 single cells in the merged ICM-EPI-PE data frame of human and *Cynomolgus*. Variation due to batch effects was adjusted using COMBAT (Johnson et al., 2007) from the R package sva. We checked the normalization status by drawing PC biplots using various subsets of clustered genes to ensure that cells did not cluster on the basis of platform or species. We checked the validation of this analysis by visualizing the selected gene expression (log TPM values) of conserved lineage markers across vertebrate blastocysts (Nakamura et al., 2016). Notably, plots show a similar expression pattern of SPIC (ICM) NANOG, POU5F1, (ICM/EPI, NODAL, GDF3 and PRDM14 (EPI), APOA1, GATA4 and COL4A1 (PE) DLX3, STS and PGF (TE) in both human and macaque.

### Cross-species normalization

Cross-species single cell datasets were normalized using the recently published *Seurat Alignment* function (Butler et al., 2018). First, we processed ICM, EPI and PE cells from either *Homo* or *Cynomolgus* using orthologous genes. We found similar expression patterns of conserved markers in the lineages of both species. We then detected variable genes in each of the datasets, using the *FindVariableGenes* function with default parameters of ‘Seurat’. We then merged the log normalized and scaled datasets from both species into a single dataset. As defined in the manual, we used all unique genes from the intersection of the two variable gene sets from *Homo* and *Cynomolgus* dataset. We used this gene set as an input to canonical correlation analysis (CCA), and alignment was performed using canonical correlation vectors (CCV) across datasets with the *AlignSubspace* function. This normalized set of cross-species data was used for downstream tSNE analysis. tSNE visualization was done of the first two dimension (tSNE1 and tSNE2). The dimension reduction method used was PCA with the *cca.aligned* parameter providing the first two dimensions.

### Cross-species trajectory analysis

As defined previously, we imported the CCA normalized cross-species Seurat object into “Monocle2” workspace. We then processed the dataset in a similar fashion for visualization of their trajectories upon providing the top 100 differentially expressed genes obtained from *differentialGeneTest* (a built-in function from “Monocle2”). These genes were treated as ordering genes for the trajectory analysis. We used our annotations of human and macaque blastocyst lineages in order to label the cells on the cross-species trajectory (Figure 2C). Note: we found a conserved cross-species trajectory of ICM to EPI and PE using top 1000 DEGs. We then lowered the number of input genes in an unsupervised manner to find the next node in the trajectory. Consequently, we observed another node bifurcating *Homo* and *Cynomolgus* EPI lineages using the top 250 most DEGs. This observation was consistent with the top 100 DEGs (Table S4).

### Cross-species markers

Cross-species markers in this study are based on orthologous genes of human and *Cynomolgus* ICM, EPI and PE lineages. Thus, we do not consider the species-specific genes that cannot, by definition, be expressed in both species (e.g. NANOGNB-ICM, ESRG-ICM/EPI, LINC00261-PE, etc.). To determine species-specific expression of orthologous genes, we use the “percent of cells expressing a given gene” criteria: (i) genes that are expressed in > 95% of the cells in a focal lineage of one species; (ii) in < 5% of cells in the same lineage of the comparator species; and (iii) are significantly upregulated in the focal lineage (*differentialGeneTest* built-in function from “Monocle2”) are considered to be cross-species markers of that focal lineage.

### Self-renewal regulatory network

We created a data frame of single-cell data for ICM, EPI, PE and TE from days E5, E6 and E7, carrying expression values (TPM) of all human genes. We used the “SCENIC” R package which depends on “GENIE3”, “AUCell” and “RcisTarget” to score the enrichment of different gene sets per single cell in unbiased manner (Aibar et al., 2017). This method is independent of the gene expression units and the normalization procedure. AUCell then uses the Area Under recovery Curve to calculate whether a critical subset of the input gene set is enriched at the top of the rankings for each cell. The AUC value that is plotted between the first two tSNEs (Figure 6E) represents the proportion of expressed genes in the signature and their relative expression values compared to the rest of the genes within the cell. Distinct gene sets that were obtained with the above procedure were stored in their respective modules/regulons. Amongst the top regulons that we obtained, we show the one which has highest enrichment score in EPI comprised of GDF3, NODAL, LEFTY1, TDGF1/CRIPTO, ESRG, TDGF1P3, IFITM1, SCGB3A2 etc shown in (Figure S4C). This module was only enriched in EPI and contained the genes that have been established as self-renewal genes. We then computed pairwise ranked Pearson correlations for the genes in all the enriched regulons. We then selected only those paired genes that show a strong correlation or anti-correlation (threshold rho > |0.80|), as shown in a heatmap (Figure S4B). In order to construct and compare the self-renewal network, we removed those genes that are not annotated in *Cynomolgous*. A network was constructed based on genes with the highest level of correlation among each other, with r > 0.80, using igraph (http://igraph.org/r/) package from R. We considered these genes as paired. Arrows show the linking between genes (links based on a preset level of preferential attachment (Barabasi-Albert model)). The direction of the arrows was manually set to illustrate the sequential order of gene expression during progression. The size of a circle represents the number of instances when the expression of a particular gene is upstream (nodes) or paired (edges). Genes in the network are markers of human EPI.

### Visualization of reads

Mapped reads from single cell transcriptomes of human embryonic development were merged for each stage, defined as EPI, PE and TE (Yan et al., 2013). Mapped reads in bam format were converted into a signal file in bedGraph format to visualize through IGV over Refseq genes (hg19) using STAR with parameters –*runMode inputAlignmentsFromBAM –outWigType bedGraph –outWigStrand Untranded*. Signals from ATAC-seq and ChIP-seq were obtained from MACS2 after running over with parameter -g hs -q 0.01 -B. The conservation track was visualized through UCSC genome browser under net/chain alignment of given non-human primates (NHPs) shown in Figure 4D and, later on, merged beneath IGV tracks.

### Detection of chimeric transcripts from RNAseq data

In order to determine chimeric transcripts, we first aligned the reads using universal aligner STAR using the parameters given above (see data processing) that can discover canonical and non-canonical splice and chimeric (fusion) sites. We retained only junctions that were identified with a minimum of 6 uniquely mapped reads. Any novel genes with resemblance to mitochondrial genes were excluded from the analysis (e.g. donor or acceptor sites mapping to the mitochondrial genome were considered grounds for exclusion). To exclude chimeras derived from repeated elements, we identified those novel transcripts that had at least 6 consecutive bps from known repeated elements (repeat specified in hg19 rmsk.gtf).

### Human embryo manipulation and microscopy analyses

Prior to the start of the project, the whole procedure was approved by local regulatory authorities and the Spanish National Embryo steering committee. Cryopreserved human embryos of the maximum quality were donated with informed consent by couples that had already have undergone an *in vitro* fertilization (IVF) cycle. All extractions/manipulations were carried out in a GMP certified facility by certified embryologist in Banco Andaluz Celulas Madre, Granada, Spain. Confocal analyses of LINE-1 ORF1p expression were analysed on a Zeiss LSM 710 confocal microscope using a previously described method (Macia et al., 2017). Antibodies for the immunostaining: Rabbit anti LINE-1 ORF1p, 1:500, a generous gift of Dr Oliver Weichenrieder (Max Planck, Germany). Secondary antibody: Alexa 488 Donkey anti Rabbit, 1:1000 (Thermo). Mouse anti H2AXγ, 1:200, clone 3F2 (Novus). Secondary antibody: Alexa 555 Donkey anti Mouse, 1:1000 (Thermo). DAPI (Thermo) was used at 1:500. To quantify the number of L1-ORF1p cytoplasmic foci in human blastocysts, every second image of a confocal stack was used at the height of the ICM. L1-ORF1p cytoplasmic foci were enhanced by applying a granulation filter with the same parameters to all images, and ORF1p cytoplasmic foci were counted manually.

## Acknowledgments

Z.I. is funded by European Research Council, ERC Advanced [ERC-2011-ADG 294742]. L.D.H. is funded by European Research Council, ERC Advanced [ERC-2014-ADG 669207]. J.L.G.P’s lab is supported by CICE-FEDER-P12-CTS-2256, Plan Nacional de I+D+I 2008-2011 and 2013-2016 (FIS-FEDER-PI14/02152), PCIN-2014-115-ERA-NET NEURON II, the European Research Council (ERC-Consolidator ERC-STG-2012-233764), by an International Early Career Scientist grant from the Howard Hughes Medical Institute (IECS-55007420), by The Wellcome Trust-University of Edinburgh Institutional Strategic Support Fund (ISFF2) and by a private donation by Ms Francisca Serrano (*Trading y Bolsa para Torpes*, Granada, Spain).

## Author contributions

The authors declare that they have no conflicts of interest. Z.I., M.S and L.D.H. designed the study and drafted the manuscript. M.S. conceived the idea, designed and performed the computational analyses. V.B. assisted with the trajectory analysis and performed SCENIC analyses. S. W. and U. M. have assisted in providing the NHP lines. G.G.S. provided the material isolated from the Callithrix ESC line CJES-001. The official provider of the *Callithrix* Embryonic stem cell line is the Central Institute for Experimental Animals (CIEA), 1430 Nogawa, Miyamae, Kawasaki 216-0001. T.J.W, J.L.C. and J.L.G.-P. carried out all the work with cryopreserved human embryos.

## Competing interest statement

The authors declare that they have no competing financial interests.

**Supplementary Information is linked to the online version of the paper.**

## Supplementary Figures

**Figure S1, related to Figure 1.**
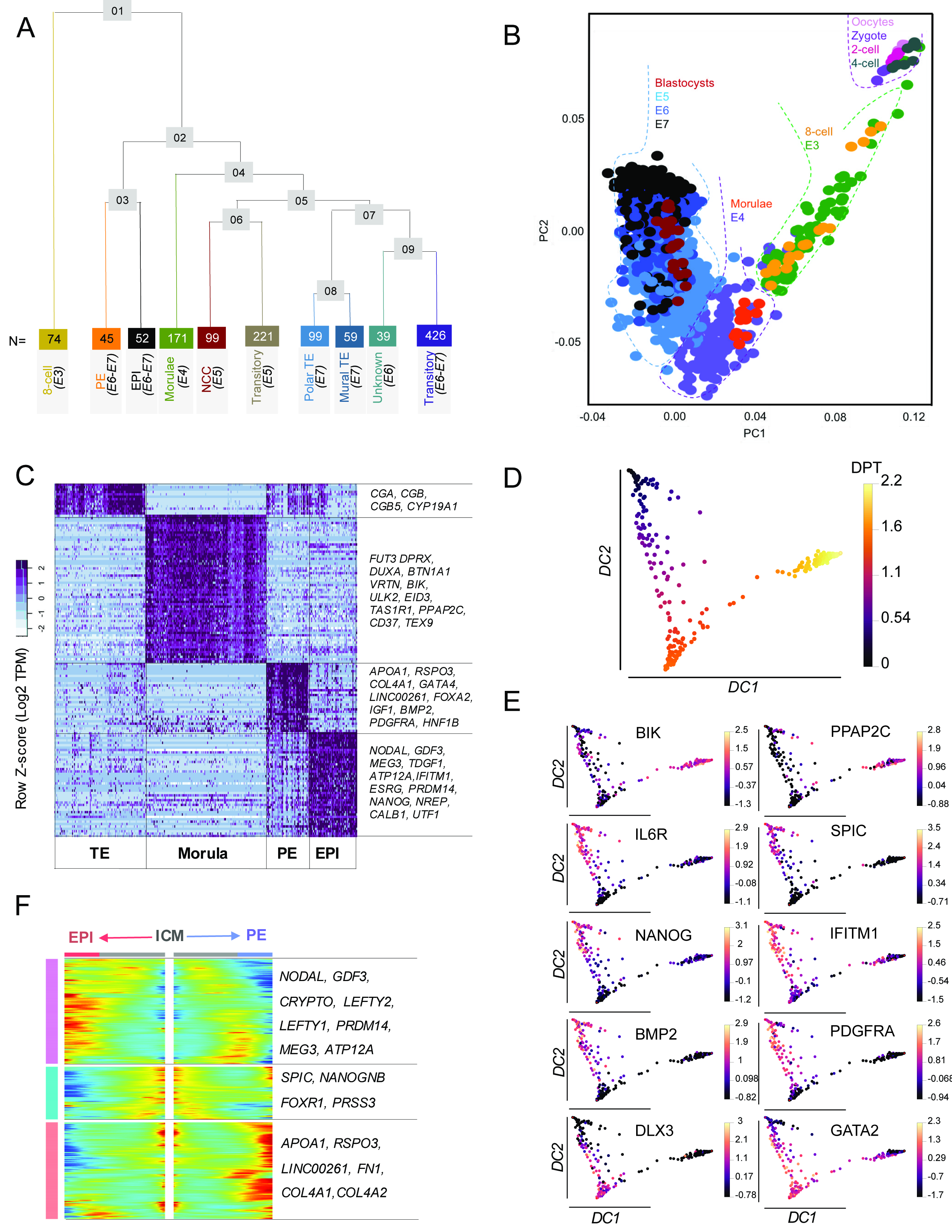
Dissecting human pre-implantation embryogenesis. A. Re-ordered phylogenetic tree of clusters shown on Figure 1A. The tree is constructed using “*BuildClustertree*” built-in function of the R package ‘*Seurat*’. Nodes are shown in grey boxes and numbers are in the order of their position on the tree. Colour codes are as in Figure 1A. Numbers in coloured boxes denote the number of cells in each representative cluster. Note: From the 1285 single cells (Petropoulos et al., 2016), the independently clustering 16 single cells were added to the transitory category from E6 and E7 (originally 410 cells). At E5, a cluster of 99 cells does not express any of the lineage markers (non-committed cells, NCCs) (dark red). B. Tracing the human embryonic development progression from zygote to blastocyst. Principal component analysis (PCA) of cross-platform 1285+114 single-cell pre-implantation transcriptome (Petropoulos et al., 2016; Yan et al., 2013) using 1,583 most variable genes (MVGs). Developmental stages defined as in (Blakeley et al., 2015; Petropoulos et al., 2016). C. Heatmap displaying the scaled expression (Log2 TPM values) of discriminative gene sets (AUC cutoff ≥ 0.90) defining cell populations of morula (n=171), EPI (n=52), PE (n=45) and TE (n=99) reported in Figures 1A and S1A. Heatmap colour scheme is based on z-score distribution from −2 (light blue) to 2 (purple). Note: The majority of the markers agrees with the previous study (Petropoulos et al., 2016), where EPI, PE and TE genes were defined as differentially expressed at defined embryonic days (E5, E6 and E7). Here we show the markers of these lineages from E3-E7. D. Diffusion pseudotime (DPT) plot between Diffusion Component 1 and 2 (DC1 and DC2) illustrating three major states of E5 along pseudotemporal ordering. Bottom: pre-TE. top: ICM. Middle right: NCCs and cells are under progression. E. The series of feature plots show the expression dynamics of individual genes in E5 blastocyst plotted on the DPT (Figure S1D). Cells coloured dark and golden-yellow are representing higher and lower expression of respective genes, respectively. ICM (e.g. IL6R and SPIC) is characterized by the progressing cells enriched in EPI (e.g. NANOG and IFITM1) and PE (e.g. BMP2 and PDGFRA). pre-TE population is identified by marker gene expression (e.g. DLX3 and GATA2). NCCs do not express any lineage markers, and marked by BIK and PPAP2C expression. F. Heatmap showing the kinetics of genes changing gradually over the trajectory of ICM differentiation to EPI or PE. Genes (row) are clustered and cells (column) are ordered according to the pseudotime progression. Genes projected in EPI are associated with self-renewal (NODAL, GDF3, LEFTY1/Y2, CRYPTO), ICM being the progenitor lineage is marked by SPIC, NANOGNB, FOXR1, whereas, PE projections are determined by APOA1, RSPO3, COL4A1/A2 and FN1.

**Figure S2, related to Figures 1 and 2.**
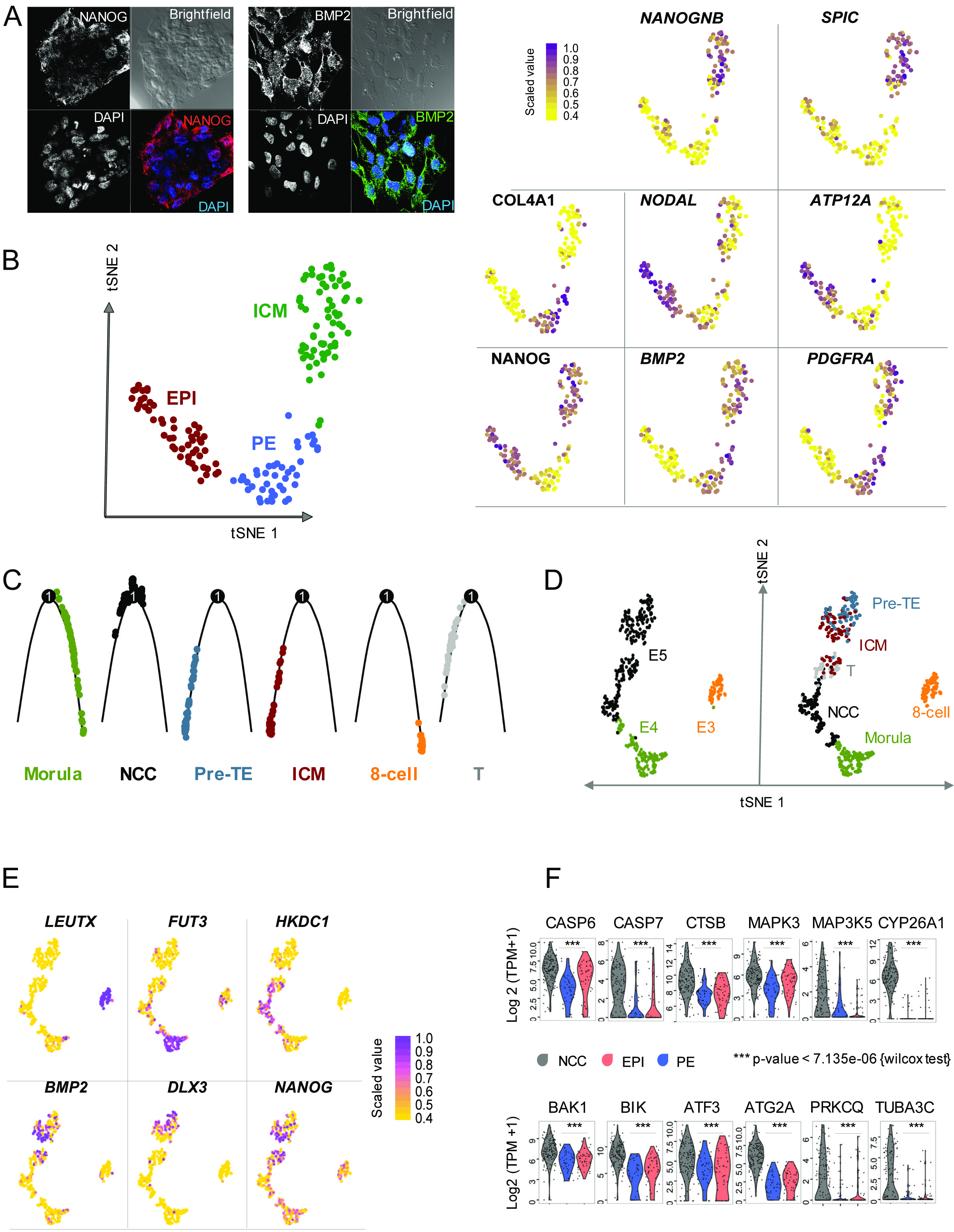
Committed and non-committed cell fates during human blastocyst formation. A. Optimisation of immunostaining using NANOG (red) and BMP2 (green) antibodies in ESC_H9 cells. B. Multiple feature plots displaying unsupervised identification of precise expression markers of ICM (NANOGBB, SPIC), ICM/EPI (NANOG, SCGB3A2), ICM/PE (BMP2, PDGFRA), EPI (NODAL, ATP12A) and PE (COL4A1, APOA1) (see also Figure 1I and Table S3). Dots in yellow denote lower whereas purple higher level of gene expression in a given single cell. Note: tSNE plot of ICM, EPI and PE (see also Figure 1I), is shown as a reference for the feature plots (bottom left). C. Monocle2” single cell trajectory analysis and ordered cells along an artificial temporal continuum using the top 1000 differentially expressed genes across the data frame of E3-E5 cell populations shown on (Figure S2D). The transcriptome from each single cell represents a pseudotime point along an artificial time vector that denotes the progression from 8-cell to blastocyst via morula. Note: The artificial time point progression agrees with the biological one. NCCs deviate on the trajectory. We show the trajectory in six facets, one for each cluster identified previously (Figure S2D). Coloured codes as on Figure S2D, right panel. D. tSNE clustering of E3-E5 cells from ~ 1000 MVGs obtained using default parameters of “Seurat” package (left panel), reveals five groups of cells. The analysis dissects three distinct populations of E5 blastocyst named as ICM, pre-TE and NCC. Transitory cells are observed between ICM and NCC. Each dot represents an individual cell plotted on first two tSNE (right panel). E. Feature plots based on previous tSNE plot (Figure S2D) demonstrating lineage-specific expression of LEUTX (8-cell marker (Jouhilahti et al., 2016; Petropoulos et al., 2016; Yan et al., 2013), FUT3 (Morula marker (Petropoulos et al., 2016; Yan et al., 2013), HKDC1 (Morula 1 and NCC marker, this study) NANOG-BMP2 (ICM marker, this study) and DLX3 (pre-TE marker (Petropoulos et al., 2016). Dots in yellow denote lower whereas purple higher level of gene expression in a given single cell. F. Multiple violin plots visualize the density and distribution of expression (Log2 TPM values) of selected genes that are upregulated in human NCC vs EPI/PE. The depicted genes are involved in regulating apoptotic pathways (KEGG: hsa04210, Gene Ontology GO:008219, GO:0012501 and GO:0006915) (Wilcoxon test, p-value < 7.135e-06).

**Figure S3, related to Figure 3.**
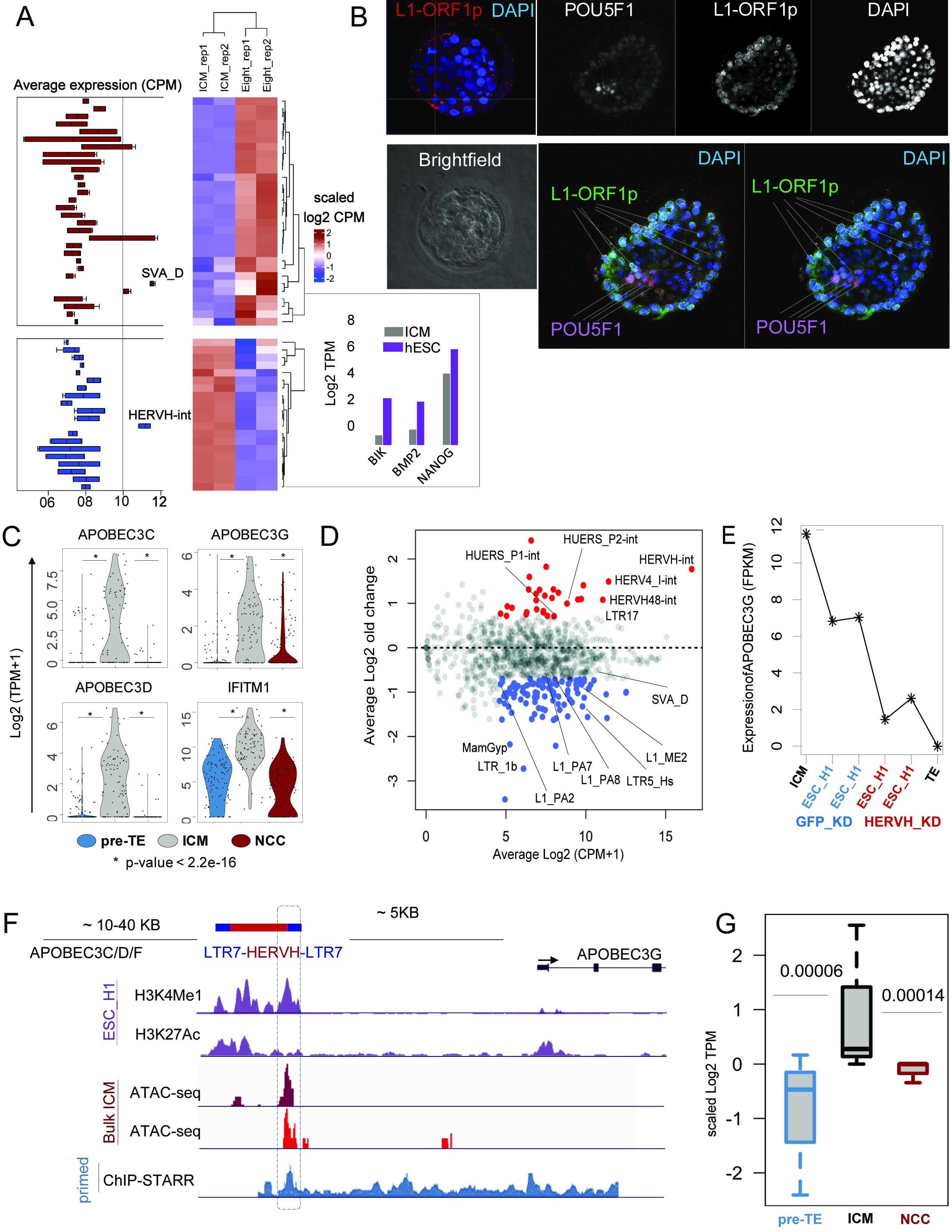
Upregulated apoptotic gene expression and Young transposable elements in non-committed cells. A. Combined boxplots and heatmaps showing distinct pattern of highly expressed TrEs families at Day 3 (8 cell) and Day 5 (bulk-ICM) of human preimplantation embryogenesis. Conventional RNAseq is not suitable to distinguish between ICM and NCC, and bulk-ICM cells express both ICM (e.g. NANOG, BMP2) and NCC (e.g. BIK) markers (inset). SVA_D/E and HERVH-int are the most abundant TrEs in the transcriptome of 8-cell and bulk-ICM, respectively, and possess an opposite dynamic of expression. B. Representative immunofluorescent staining against L1-ORF1p (red) in the human blastocyst at E5 (DAPI, blue). Note that L1-ORF1p expression preferentially accumulates in the cytoplasm of pre-TE and blastocoel cells (left panel). A representative POU5F1/OCT4, L1-ORF1p and DAPI co-staining performed on E5-Late cells shows antagonistic expression of POU5F1/OCT4 and L1-ORF1p in the embryo (right panels). Bright field images (upper panels). Merged immunofluorescent/DAPI image of the stained blastocyst are shown (POU5F1/OCT4, purple; L1-ORF1p, green). For numerical analysis of the antagonistic staining of POU5F1/OCT4 and L1-ORF1p see Figure 3C. C. Violin plots visualize the density and expressional dynamics of APOBEC3C/D/G and IFITM1, implicated in host-defence against retroelements and viruses, in non-committed (NCC) vs committed cells of pre-TE and ICM (E5, human blastocyst). Note: the transcription of the depicted genes mark ICM. D. MA plot displaying the comparison of average difference of normalized expression (CPM) of various transposable element (TrE) families in knockdown HERVH vs control cells (Lu et al., 2014). Y-axis, Log-fold change GFPcontrol-KD (n=2) vs HERVH_KD (n=2) in ESCs_H1; X-axis, average expression of TrE families in the same dataset. Dots represent TrE families downregulated (blue) or upregulated (red) (Log2 average CPM > 2 and average difference > 2) in HERVH-KD/ESC_H1. E. Line plot showing the expression of APOBEC3G in ICM (positive control), GFPcontrol-KD (two replicates), HERVH-KD (two replicates) in ESC_H1 and TE (negative control). F. HERVH as a functional enhancer of APOBEC3G. Genome browser view showing H3K4Me1, H3K27Ac, ChIP-STARR-seq (ESC_H1), ATAC-seq (two replicates of freshly isolated human bulk-ICM)) signals at the APOBEC3G locus, including the upstream full-length LTR7-HERVH-LTR7 in human PSCs. Framed region highlights the overlapping peaks at the HERVH. G. Upregulated gene expression neighbouring HERVH loci (10 KB window) in ICM. Boxplot represents the distribution of relative average difference (at Log2 scale) of gene expression at HERVH neighbours. Note: Cells are pooled together, scaled and averaged for the analysis. Only genes, neighbouring HERVH (10KB window) and expressed at least in 10% of cells are used for the analysis. Upregulated gene expression neighbouring HERVH (HERVH target) was specifically observed in ICM but not in NCC and pre-TE transcriptomes.

**Figure S4, related to Figure 4.**
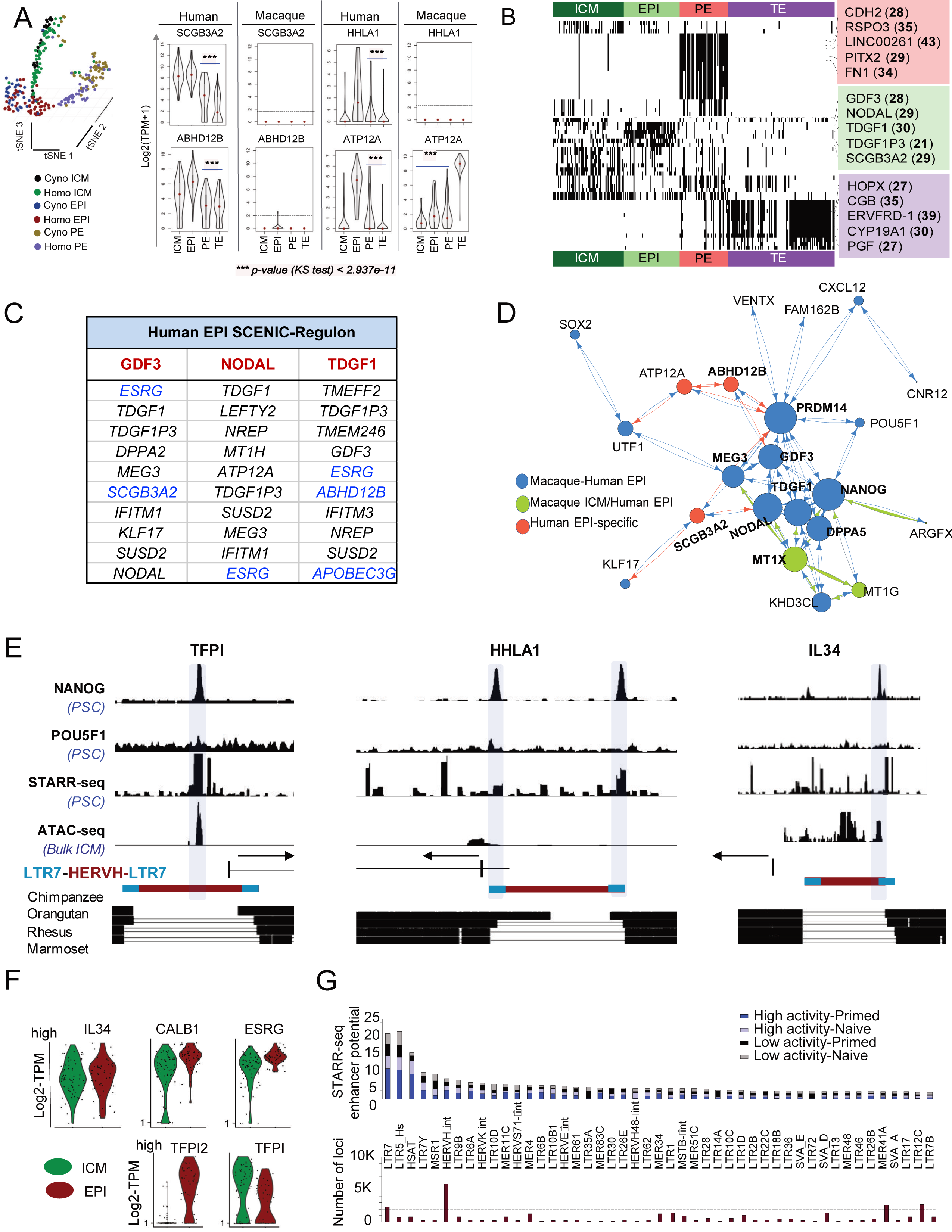
Functional divergence of pluripotent cell types in the primate blastocyst. A. Transcriptional divergence of the pluripotent EPI between *Cynomolgus* and *Homo* by analysing single cell transcriptomes. Nonlinear 3-D tSNE visualization of merged and normalized transcriptomes from human (169) and macaque (118) ICM, EPI and PE lineages. Every dot represents a single cell. The annotation is based on the clusters depicted from Figure 4B (left panel). Species-specific gene expression marks distinct stages of early human development. Violin plots visualize the density and distribution of gene expression (Log2 TPM values) of selected orthologous genes in the human vs macaque blastocyst lineages). Note: Selected HERVH-remodelled genes (e.g. SCGB3A2, ABHD12B and HHLA1) are specifically marking human pluripotent lineages. Expression of ATP12A (not HERVH-dependent) is shifted from macaque TE to human EPI (right panel). B. Barcoding human blastocyst lineages with regulons (gene sets that are co-expressed in single cell clusters). Heatmap on the binary activity matrices after applying SCENIC. The activity status of the regulon in a particular cell: active (black) or not active (white). Rows are clustered using Pearson’s coefficient. Master regulators are in boxes, colour matched with the lineage they regulate. Colour codes for the blastocyst lineages as in Figure 6E. Number of genes enlisted in a respective regulon are in brackets. The full list of genes is provided in Table S7. AUC values (quantified by AUCell) are further transformed into a binary activity matrix as suggested by SCENIC (Aibar et al., 2017). C. Evolution of the transcriptional network regulating self-renewal between *Cynomolgus* and *Homo* by analysing single cell transcriptomes. Note: Due to its low cell-to-cell variation, it is possible to obtain gene pairs in the transcriptome of the human EPI. Only strongly correlated pairs (Pearson’s coefficient > 80%), and genes annotated in both species are considered for the analysis. For each pair, nodes and edges are decided on their expressional dynamics on pseudotime scale in the EPI cluster. The size of the nodes is proportional to the number of components the representative node is paired with in the network (see methods). Colour codes: gene expression is shifted from macaque ICM to human EPI (green); expressed in both human and macaque EPI (blue); expressed in human EPI only (red). D. Top 10 key genes of the NODAL, GDF3 and TGDF1/CRIPTO regulons of human EPI (see also Figure S4B and Table S7. Human specific HERVH-target genes are in blue. E. HERVH as a functional enhancer of TFPI, HHLA1 and IL34 in human pluripotent stem cells (PSCs). Genome browser view showing NANOG (ESC_H9), POU5F1 (ESC_H9), ChIP-STARR-seq (hPSC), ATAC-seq (bulk-ICM), signals at the depicted genomic loci, including the upstream full-length LTR7-HERVH-LTR7 in human PSCs. Shading highlights the overlapping peaks. Arrows show the transcription start sites (TSS) and the direction of the transcription of the genes. Note that HERVH overrides annotated gene regulation and modulates expression. Bottom panels show phylogenetic conservation status of the loci, the presence (thick line) and the absence (narrow line) of the human sequence compared to the *Chimpanzee*, *Orangutan*, *Rhesus* and *Marmoset* assemblies. F. Violin plots visualize the density and distribution of expression of depicted HERVH-target genes (e.g. IL34, CALB1, ESRG, TFPI2 and TFPI) in the human EPI (dark red) and ICM (green). Note the differential/exclusive gene expression of the TFPI paralogues in the pluripotent cell types of ICM and EPI. Each dot represents an individual cell. G. Relative functional enhancer potential of various TrE-derived sequences (top 48) determined by ChiP-STARR-seq (GSE99631) in primed and naive PSCs (stacked bar plots). The lower panel displays the total numbers of the particular TrE-derived sequence loci in the human genome (corresponds to the upper panel).

**Figure S5, related to Figure 5.**
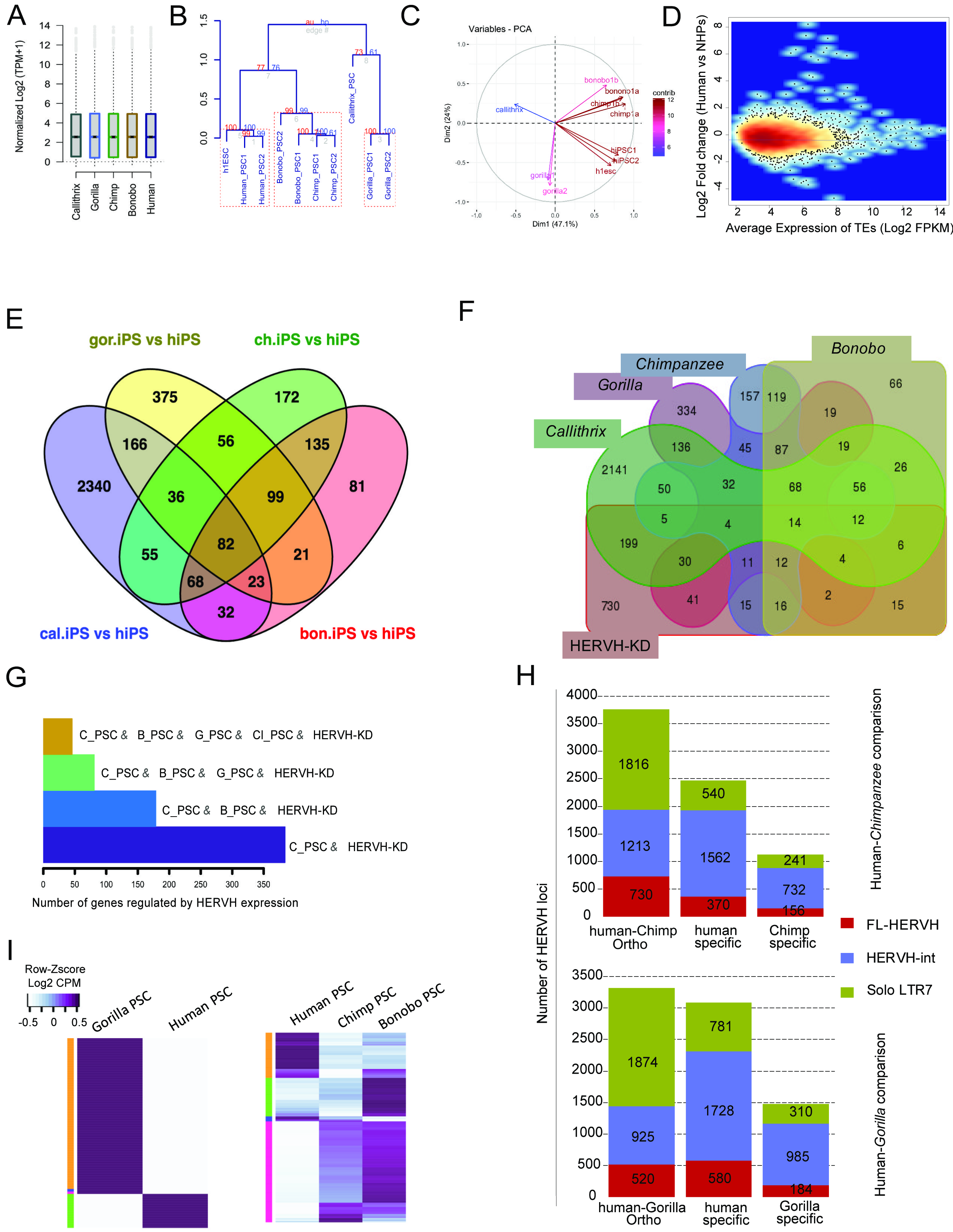
The robust divergence of pluripotency regulation in primates is HERVH-enforced. A. Quality control. Boxplots show the normalized global expression estimates of RNAseq datasets from various primate pluripotent stem cells (PSCs) analysed in this study (cross-species mapped). B. Quality control. Correlation of the transcriptomes of the primate samples used in the study. The dendrogram showing hierarchical clustering using ranked correlation and complete linkage method on scaled expression data (1000 bootstraps). Height of the dendrograms represent the Euclidian distance of the dissimilarity matrix, numbers in red and blue indicate au and bp values from bootstrapping. Note the species-specific clusters. C. Quality control. Projected PCA on the significantly expressed TrEs (Log2 FPKM > 1) following cross-species mapping of high quality RNAseq reads (MAPQ > 30). Batch effect was further removed by “combat” function of the “sva” package. This plot shows the normalized species-specific pattern of TrEs in PSCs. D. Quality control. MA-plots demonstrates the normalization of TrE expression in cross-species comparisons. This differential expression plot is drawn between Log2 fold change of human/NHP PSCs vs average expression TrEs of all analysed samples. E. Venn diagram displays the divergence of PSC transcriptomes in primates (e.g. human, *Bonobo*, *Chimpanzee*, *Gorilla* and *Callithrix*). The numbers in the Venn diagram denote differentially expressed genes (DEGs) (FDR<0.05 and fold-change |2|). RNAseq data of various non-human primate (NHP) PSCs were compared to human PSCs. Only cross-species reads mappable to both genomes were considered for the analysis. Gene expression was calculated on the human genome, using the human gene models. F. The impact of HERVH-mediated regulation on the evolution of primate pluripotency. As in Figure S5E, but also including differentially expressed genes (DEGs) upon HERVH knockdown (HERVH-KD in ESC_H1 vs GFPcontrol-KD in ESC_H1). G. Barplot showing the number of DEGs affected by HERVH expression. As in Figure S5C, but also including DEGs genes upon HERVH knockdown (HERVH-KD in ESC_H1 vs GFPcontrol-KD in ESC_H1). H. Stacked barplot showing the number of HERVH orthologous or species-specific in human-chimp (upper panel) or human-gorilla (lower panel) genome-wide comparisons. FL, Full-length. I. Heatmaps display the loss and gain of expression of orthologous HERVH loci between human vs gorilla (left panel) and human vs chimp/Bonobo PSCs. Only RNAseq reads, mappable to both human and NHP reference genomes were considered.

**Figure S6, related to Figure 6.**
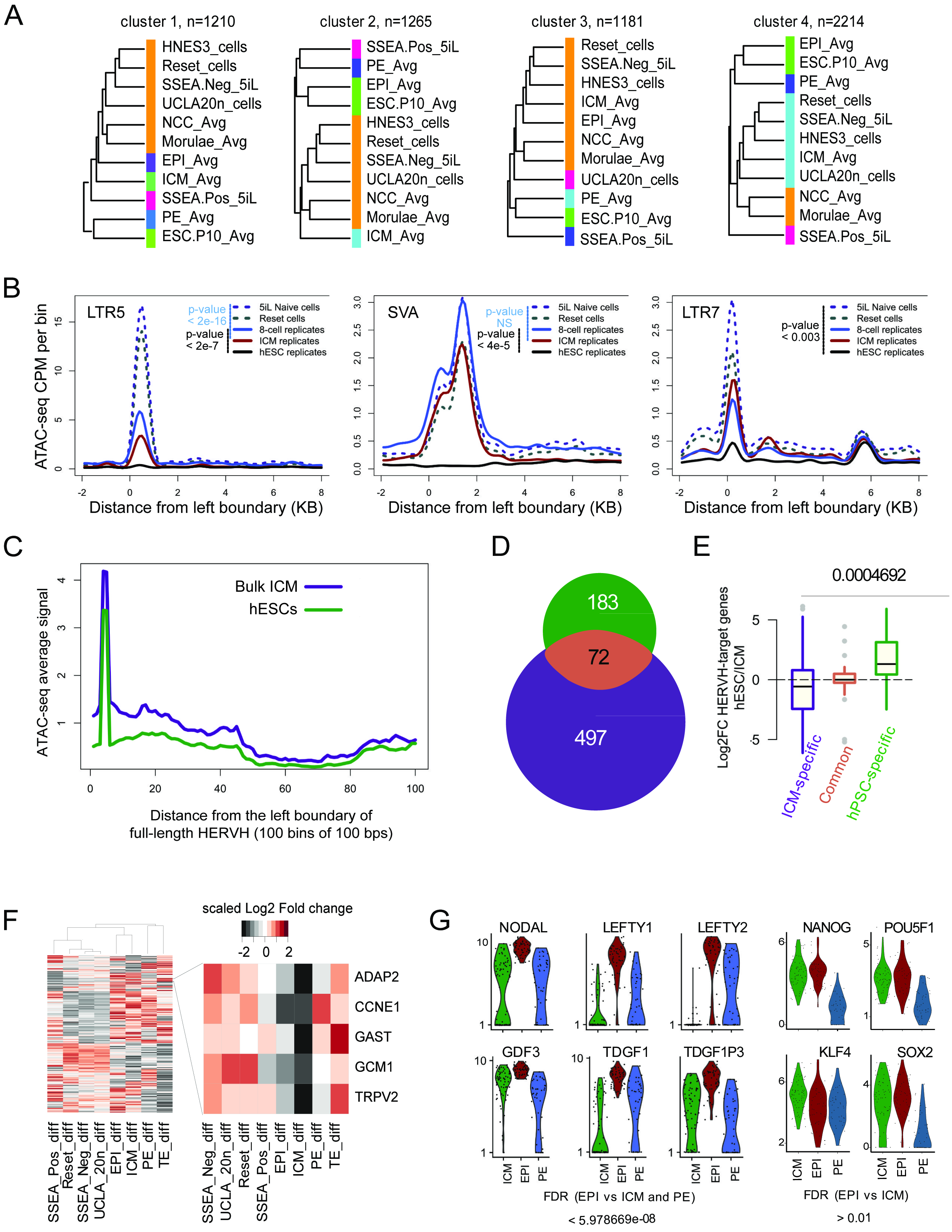
Lessons for *in vitro* models of early embryogenesis. A. Dendrograms show the four clusters obtained from transcriptome-wide clustering of samples based on Euclidian distance of dissimilarity matrix using 1210, 1265, 1181 and 2214 genes from cluster 1 to cluster 4. Reset/ESC_H9 (Takashima et al., 2014), 5iL_SSEA_Neg/UCLA1_primed, UCLA_20n/UCLA_20n_primed, 5iL_SSEA4_Pos/UCLA1_primed (Pastor et al., 2016) and Chan_3iL/ESC_H1s (Chan et al., 2013). Note: Depending on the input gene sets, naïve cultures cluster with NCC, morula, ICM or EPI. B. ATAC-seq coverage plots show the signals around the transcription start sites (TSSs) located at the left boundaries of SVA, LTR5 and LTR7 loci. X-axis, upstream 2 KB and downstream 8 KB regions from the left boundaries divided into 100 bins, each comprising 100 bps; Y-axis, normalized ATAC-seq signal (per million per 100 bp bin). Samples: Replicates (n=2) of human 8-cell stage embryo, human bulk-ICM and hESC_H1 (solid lines), Reset-naive, 5iL-naive cultures (dashed lines). The P values are calculated with Wilcoxon method and further adjusted by BH method. C. ATAC-seq coverage plot shows the signals around the transcription start sites (TSSs) located at the left boundaries of full-length LTR7/HERVH loci. X-axis, 100 bps bins up to 10 KB; Y-axis is normalized ATAC-seq signal of in human bulk-ICM (purple) and hESCs (green). D. Venn diagram of ATAC-seq enriched regions (FPKM > 2) over HERVH between human bulk-ICM (497) and cultured ESC_H1 cells (183) with an overlap (72) (colour codes as on Figure 6B). E. Boxplot shows the relative expression of HERVH-target genes in cultured hESCs and bulk-ICM (specific vs common). Note that HERVH-target genes shared between bulk ICM and cultured hESCs are not expressed differentially at significant level. F. Heatmap showing the comparison of expressional changes as fold-change of naïve vs primed cells with the fold-change of pairwise human and macaque blastocyst stages (ICM, EPI, PE and TE). Row-wise z-score of Log2-fold change expression of most variable genes (MVGs) (n=948). The clustered dendrogram represents Spearman’s correlation and Euclidian distance. Note the contrasting pattern of gene expression between *in vitro* naïve cultures and human *in vivo* pluripotent states (ICM and EPI) compared with their counter samples. The zoom-in details expressional changes of five genes, whose expression is shifted between macaque and human (e.g. mark pluripotent states in macaque and TE in human). The five genes also represent genes that are upregulated in naïve *in vitro* cultures vs their primed counterparts. Note: Since, this comparison involved single vs bulk RNA-seq datasets, we had pooled all single cells belonging to respective lineage into one file and treated as a sample for comparison. G. Violin plots illustrate gene expression distribution of selected genes associated with pluripotency (e.g. NANOG, POU5F1, SOX2, KLF4) and self-renewal (e.g. NODAL, GDF3, TDGF1, TDGF1P3, LEFTY1/2) in human ICM, EPI and PE (p-value < 0.00005) (colour codes as in Figure 1J).

## Supplementary Tables

**Table S1**

Transcriptional markers of morula and blastocyst lineages

**Table S2**

235 marker genes as molecular signatures of the individual clusters at E5 blastocyst (ICM, NCC and Pre-TE)

**Table S3**

Expression markers of ICM, ICM/EPI, ICM/PE, EPI and PE in human blastocyst

**Table S4**

Phylogenetically conserved expression markers of ICM, ICM/EPI, ICM/PE, EPI and PE in primate blastocyst from orthologous set of genes

**Table S5**

Species specific expression markers of ICM, ICM/EPI, ICM/PE, EPI and PE in primate blastocyst from orthologous set of genes

**Table S6**

Differentially expressed genes (DEGs) between human and non-human primate pluripotent stem cells

**Table S7**

Top five regulons of human blastocyst lineages

**Movie S1**

Expression dynamics of L1-ORF1 (red) by immunofluorescence staining during the formation of the blastocyst. Note that L1_HS_ORF1p accumulates in the cytoplasm of Pre-TE and in the blastocoel cells (DAPI, blue). See also Figure S3A.

